# Identifying the metacommunity scale reconciles deterministic and stochastic assembly in a natural mainland–island microbial system

**DOI:** 10.64898/2026.11.15.766812

**Authors:** Javier Cuadrado Corz, Josep Ramoneda, Diego Alcón, Thomas Bell, Alberto Pascual-García

**Affiliations:** Department of Life Sciences, Imperial College London, Silwood Park Campus, Ascot, UK; Centre for Cancer Biomarkers and Biotherapeutics, Barts Cancer Institute, Queen Mary University of London, London, UK; Centre for Advanced Studies of Blanes (CSIC), Blanes, Spain; Spanish National Centre for Biotechnology (CSIC), Madrid, Spain

## Abstract

Disentangling the deterministic and stochastic processes that shape microbial communities remains a central challenge in microbial ecology. The framework proposed by Vellend—selection, drift, dispersal and speciation—offers a natural conceptual scaffold, but testing these processes in natural communities is technically demanding, and analyses are often narrowed to a single process, which can lead to seemingly contradictory conclusions. Here we illustrate the promise and the pitfalls of a comprehensive application of this framework by re-analysing a temporally resolved dataset of microbial communities inhabiting water-filled tree-holes at the base of a single beech tree, a natural mainland–island system for which both neutral assembly and strong selection have been reported. We show that these views are compatible once unsupervised compositional clustering is used to identify the scale at which assembly processes should be tested. Treating each compositional cluster as a set of local communities that receive immigrants from a distinct metacommunity, we applied neutral, phylogenetic and null-model tests of selection and dispersal, metagenomic prediction and association inference. Selection operated chiefly as environmental filtering at the metacommunity level, associated with tree-hole size and shape, whereas local communities were statistically indistinguishable from stochastic assembly. Variable selection and dispersal limitation acted between metacommunities, whereas homogenizing dispersal prevailed within them. Association networks recapitulated metacommunity structure, indicating that co-occurrence largely reflects shared environment rather than interactions. In summary, performing some analyses while excluding others leads to misleading conclusions: comprehensive analyses and identifying the intrinsic scale in which selection may operate are needed to move beyond the deterministic–stochastic dichotomy.

## Introduction

Our ability to intervene in, or even control, natural microbial communities requires understanding the main mechanisms shaping their composition and functioning. This is a difficult task, given the vast diversity of most natural communities and the variety of pressures influencing them, from abiotic factors (e.g. pH, temperature) and environmental fluctuations (e.g. rain, seasons) to interactions among microbes or with their hosts. Our limited understanding of these processes probably contributes to the limited success of targeted interventions in natural environments (see e.g. [1, 2]).

The factors influencing the assembly of microbial communities have often been presented as mutually exclusive forces, a paradigmatic example being the debate on the relative importance of stochastic and niche-based processes. We use the term niche-based processes in the broad community-assembly sense, to refer to deterministic processes including abiotic environmental filtering and biotic interactions. Perhaps because the first attempts to address this question in macroscopic ecology originated in intense controversies, such as the one that followed Simberloff’s attempts to test the assembly rules proposed by Diamond (summarized in [3]), the inability to reject a null or neutral model [4] has often been presented as a rejection of selection. This inference is not warranted, since species with different traits can generate species abundance distributions in the statistically neutral regime [5], and environmental gradients and fluctuations may further obscure signatures of selection [6]. Deterministic and stochastic processes may therefore operate simultaneously, jointly producing the observed communities.

A step towards integrating these processes was the framework proposed by Vellend [7], who transferred the classification of processes of the Modern Evolutionary Synthesis [8] to Community Ecology and, incorporating ecological interactions as a selective force, concluded that four categories —selection, drift, speciation, and dispersal— suffice as a conceptual framework. In natural microbial communities, however, testing these processes requires specialized software to handle large numbers of taxa and artefacts such as the compositional nature of sequencing data [9], and computational tools often focus on specific processes, with few exceptions such as the pipeline proposed by Dini-Andreote and Stegen [10]. Moreover, different processes may operate at different spatial and temporal scales [11], and how to investigate whether a system has an intrinsic scale remains an open question in microbial ecology [12]. This is particularly difficult for microbes: in macroscopic systems, the local communities that share a metacommunity can often be delimited from observed movements of individuals, whereas neighbouring microbial habitats may be effectively isolated, or may receive propagules from different source pools. The scale at which stochastic and deterministic processes should be tested must therefore be inferred from the data. In the absence of comprehensive pipelines, analyses are often restricted to the processes that the researcher perceives as most relevant. Here we argue that the processes proposed by Vellend can be explored comprehensively with well-established analyses, and highlight the interpretative risks that arise when the analytical strategy is not sufficiently comprehensive. Our proposal centres on the search for clusters of compositionally similar communities, potentially subject to similar selection pressures. Their identification may reveal the intrinsic scale of the system, especially useful when appropriate metadata (e.g. spatial or temporal information) is available. We show that these clusters can be operationally interpreted as sets of local communities receiving immigration from distinct metacommunities, which enables a multi-scale description of neutral assembly, and we combine them with null models of selection, dispersal and drift, metagenomic predictions and association inference. We present the results in a different order, intentionally, to highlight the risks of an incomplete analysis.

We applied this strategy to a natural system well suited for an analogy with the classical mainland–island framework of biogeography [13]: microbial communities sampled from the interstitial water at the bottom of water-filled ponds (phytotelmata) at the base of beech trees. These ‘tree-holes’ may be regarded as ‘islands’ colonized through abiotic (e.g. rain, wind) and biotic (e.g. insects, falling leaves) transport processes which, taken together, may be abstracted as the ‘mainland’. Neutral assembly was reported in tree-hole bacterial communities [14, 15, 16], with differences between communities attributed to immigration rates determined by tree-hole surface area [16]. Conversely, strong selection was reported in a large tree-hole survey, in which communities formed compositional clusters with differentiated metagenomic and functional signatures [17]. Some of these signatures persisted after cryopreservation and repeated regrowth in the same standardized medium [18], suggesting that distinct native environments had selected for distinct and robust compositional types of communities.

To reconcile these views, we re-analysed a dataset of 10 tree-holes from a single beech tree sampled monthly over one year, together with environmental measurements (Fig. 1A and B). We found different stochastic and deterministic processes operating at the community and metacommunity levels (Fig. 1C): metacommunities were subject to variable selection, mostly associated with tree-hole size and shape, and local communities assembled stochastically from them, whereas immigration rates were at most weakly associated with tree-hole shape. Restricting the analysis to some tests leads to misleading conclusions, such as overemphasizing the role of dispersal.

**Figure 1.**
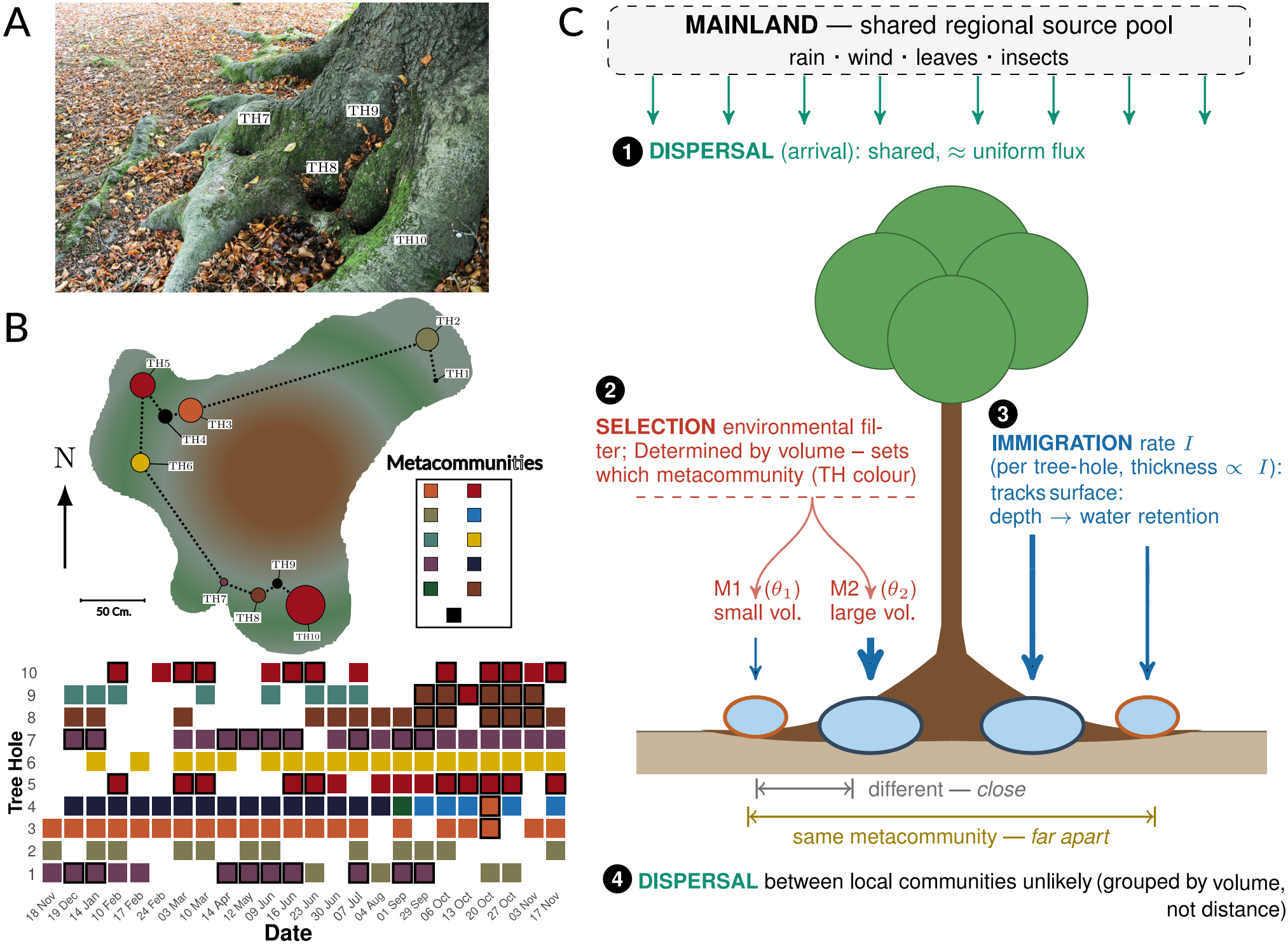
System description and conceptual framework. (A) Samples were taken from a beech tree located in Silwood Park (Ascot, United Kingdom) over the course of a year. The photograph shows some of the hollows in the tree from which samples were taken, located on the south-facing side. (B) Illustration of the base of the tree as it would appear from above. The tree cavities are shown as circles whose area is proportional to their volume. The colours represent the metacommunities to which their samples belong; three of the tree cavities have samples in several metacommunities, as shown in the inset. The matrix underneath shows the correspondence between tree-holes and metacommunities throughout time. Samples from different tree holes in the same metacommunity and date have thicker borders. (C) Conceptual framework illustrating the interaction between stochastic processes and selection, anticipating key results: selection and dispersal limitation act between metacommunities, whereas stochastic processes prevail within them (see Fig. 4A).

## Methods

### Bioinformatics pipeline

Raw sequencing data were retrieved from the authors of Ref. [16], demultiplexed with idemp (https://github.com/yhwu/idemp) and processed with DADA2 v1.32.0 [19], after trimming primers 515F [20] and 806RB [21]

with cutadapt v4.6 [22]. ASVs were inferred with sample pooling, chimeric sequences were removed (98.3% of the reads were retained), and taxonomy was assigned with the naive Bayesian classifier [23] against SILVA v138.1 [24]. After removing chloroplasts, mitochondria, negative controls, samples with 5000 reads or fewer, and ASVs with fewer than 100 reads or present in fewer than 10 samples, 3537 ASVs in 164 samples remained (details in Suppl. Methods).

### Test of neutrality

We used the multi-site neutral model of Harris *et al*. [25], a Bayesian approach that fits the matrix of ASV abundances of a set of samples (all samples, or the samples of one compositional cluster) to a Hierarchical Dirichlet Process with speciation parameter *θ*, metacommunity distribution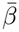 and immigration rates *I*_*i*_, one per sample. For each posterior sample of the parameters, a synthetic matrix with the same number of samples and abundances is simulated, and its metacommunity distribution is re-inferred (see Suppl. Methods). Neutrality is assessed with the proportion of posterior samples in which the log-likelihood of the observed data exceeds that of the synthetic data, an empirical (pseudo) p-value; neutrality was rejected if *p* < 0.001. In the local test,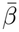 is unconstrained; in the full test, it must be consistent with a neutral metacommunity. We generated 50000 posterior realizations, discarding 25000 as burn-in and retaining 2500.

### Inference of metacommunities

Samples were classified with the Dirichlet multinomial mixture of Holmes *et al*. [26], which assumes that each community is drawn from a multinomial distribution whose parameters follow a Dirichlet prior, the conjugate prior of the multinomial. Allowing *k* subsets of samples, each with a different Dirichlet parameter vector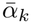, the model fits a mixture of Dirichlet distributions, and each vector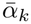 can be interpreted as a different metacommunity from which a subset of samples was drawn (see Suppl. Methods). The number of components (clusters) was the one maximizing the model evidence, which penalizes additional components [26]; each cluster defines a metacommunity (M1–M10). We used the software provided in the original article.

### Environmental drivers of composition and of neutral-model parameters

ASV counts were transformed with the centred log-ratio (CLR; pseudocount of one), and the resulting Aitchison distances were ordinated with a PCoA [27, 9]. Environmental variables were fitted onto the ordination with envfit (`vegan` [28]; 999 permutations), after log-transforming depth, surface area, volume and the depth-to-surface ratio, decomposing orientation into northness and eastness, and removing mean air and soil temperatures because of their collinearity with the maximum and minimum air temperatures (variance inflation factor *>* 10). Morphology was summarised as 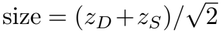 and 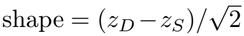, where *z*_*D*_ and *z*_*S*_ are log depth and log surface area standardised across the 10 tree-holes (Suppl. Note 1). The variance explained by each set of variables was quantified with a redundancy analysis (RDA) and the adjusted *R*^2^ [29, 30]. Morphological variables were tested by permuting morphology among the 10 tree-holes (9999 permutations), correcting for the selection of the best of 30 combinations of morphological variables with a max-statistic [31]; size and shape together were specified a priori. Time-varying variables were tested with a partial RDA conditioned on tree-hole identity, permuting samples within tree-holes, and the variance was partitioned between morphology and time-varying variables. The same procedure was applied to the log immigration rates, and the speciation rates were correlated (Spearman) with the mean of each variable per metacommunity, correcting for multiple testing [32].

### Test of selection and dispersal

Following Refs. [10, 33], we computed, for each pair of communities *k* and *l*, the weighted *β*-Mean Nearest Taxon Distance,

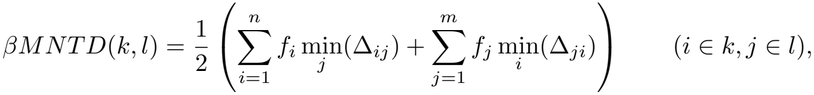

where *f*_*i*_ is the relative abundance of ASV *i* and min_*j*_(Δ_*ij*_) is the minimum phylogenetic distance between ASV *i* in community *k* and all ASVs in community *l*. A *βMNTD* significantly lower than the null expectation is expected when the environment constrains composition, and a significantly higher value when different conditions lead to divergent compositions [10]. As a null model, we shuffled the ASVs across the tips of the phylogenetic tree (999 realizations) and computed the *β*-Nearest Taxon Index [33],

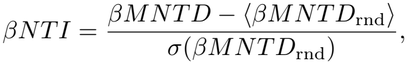

considering pairs with abs(*βN TI*) > 2 significant. For non-significant pairs we computed the Raup–Crick dissimilarity (*RC*), comparing the observed number of shared species with a null model preserving community richness and species incidence; *RC* < 0 indicates homogenizing dispersal, *RC* > 0 dispersal limitation, and values near 0 drift. Computations were conducted with the function bNTIn.p of the R package iCAMP v1.5.12 [34] with 1000 randomizations (details in Suppl. Methods).

### Metagenomic predictions

Functional profiles of the ASVs were estimated with PICRUSt 2 (SC database [35, 36]) with default parameters (median NSTI 0.13, indicating fair prediction accuracy) and mapped to KEGG pathways. The pathways found significant in this system in previous work [17] were compared between metacommunities with a Wilcoxon rank-sum test with AlDEx 2 [37] as implemented in `ggpicrust` [38], retaining features with an adjusted p-value lower than 0.05 [review criteria with last code].

### Inference of associations

A correlation network was inferred independently for each metacommunity with `SparCC` [39], as implemented in `FastSpar` [40], retaining correlations with a bootstrap pseudo *p*-value < 0.001 (1000 permutations) and |*r*| ≥ 0.3. Network similarity was quantified with the Szymkiewicz-Simpson overlap coefficient, the proportion of shared edges relative to the size of the smaller network. In M7, ASVs characteristic of each tree-hole were identified with an Indicator Value analysis (`indval` function of the `labdsv` R package [41]; strict indicators, 1000 permutations, pseudo *p* ≤0.05), complemented with ASVs showing |ΔAbundance| *>* 0.05 or |ΔRank| *>* 1000 between tree-holes (Suppl. Fig. 1), and guilds of ASVs sharing neighbours were identified with `functionInk` [42] (details in Suppl. Methods).

## Results

### Evidence of both niche effects and neutral assembly

The analysis of the 10 tree-holes from a single beech tree sampled monthly over one year revealed that tree-hole identity was a strong determinant of community composition: different tree-holes showed different taxonomic profiles (Fig. 2A and Suppl. Fig. 2; ANOSIM *R* = 0.89; *p* < 10^−3^), and communities from the same tree-hole grouped together in a PCoA ordination (Fig. 2B). In contrast, sampling time had little influence (ANOSIM, *R* = −0.034; *n*.*s*.). Because samples from the same tree-hole are not statistically independent, we treated the tree-hole as the relevant unit in subsequent analyses.

**Figure 2.**
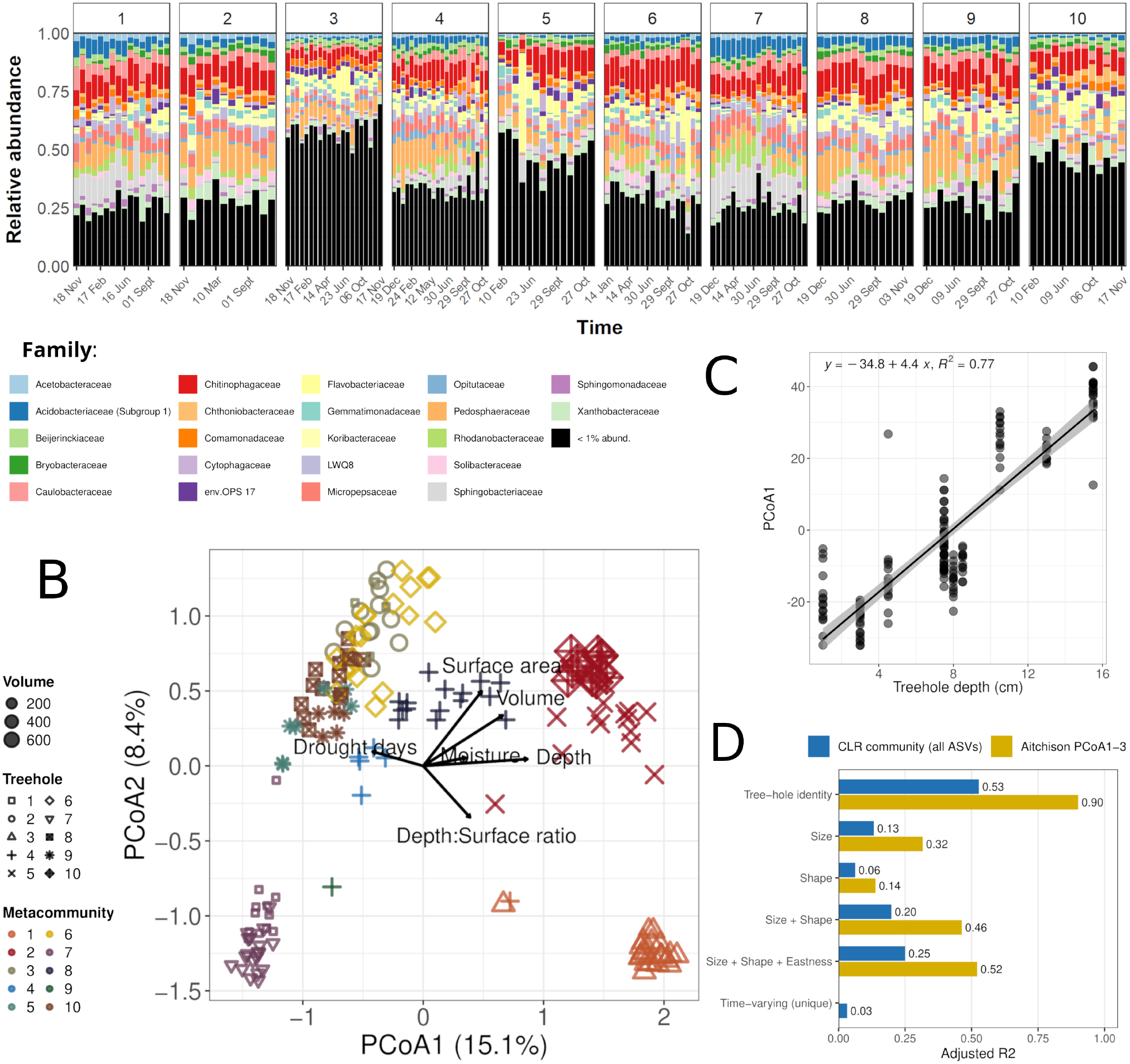
Tree-hole size and shape influence community composition. (A) Bar plots representing the ASV composition of each tree-hole (boxes) ordered by sampling time. Families representing *>* 1% relative abundances are highlighted. Most of them are apparent in all tree-holes, with different relative abundances. (B) PCoA ordination of Aitchison distances, with shapes representing the tree-hole, the size of the dots the tree-hole volume (*cm*^3^), and colours the metacommunities (see next section). Arrows represent the environmental variables significantly associated with the ordination (envfit, *P* ≤0.05), scaled by their correlation with it. (C) PCoA1 as a function of tree-hole depth, the variable most strongly associated with the ordination; the line is a least-squares fit across samples, shown for illustration (the test is reported in D). (D) Compositional variance explained (adjusted *R*^2^) by tree-hole identity, tree-hole size, shape, size and shape together, the best combination of morphological variables (size, shape and eastness) and the time-varying variables (fraction not shared with morphology), for the whole composition (all ASVs) and for the first three PCoA axes. Size and shape together were significant when morphology was permuted among tree-holes (*P* = 0.016 and *P* = 0.007, respectively; see Methods).

We investigated which environmental (e.g. temperature, humidity) or physical (e.g. surface, volume) variables were associated with this structure by fitting the measured variables onto the PCoA ordination (see Methods and Suppl. Table 1). Tree-hole depth was the variable most strongly associated with the first two axes (*r*^2^ = 0.78, *p* < 10^−3^), followed by volume (*r*^2^ = 0.49) and surface area (*r*^2^ = 0.39), and it accounted for 77% of the variation of PCoA1 (Fig. 2C; Suppl. Table 2). Among the time-varying variables, only the number of drought days (*r*^2^ = 0.21) and moisture (*r*^2^ = 0.13) were significant. Since volume and the depth-to-surface ratio are a function of depth and surface area, we searched for two uncorrelated variables: tree-hole size, which closely matched log volume (*r* = 0.998), and shape, which closely matched the log depth-to-surface ratio (*r* = 0.95) (Suppl. Fig. 3). Depth combines both dimensions, which explains why it was the best single predictor: PCoA1 was mainly associated with size, and PCoA3 with shape (Suppl. Figs.4 and 5). Because morphology is constant within each tree-hole, the effective sample size for these variables is the number of tree-holes (10) rather than the number of samples. We therefore tested them in the following by permuting the morphology among tree-holes (see Methods). Tree-hole identity explained 53% of the compositional variance when all ASVs were considered (redundancy analysis, RDA, see Methods), the upper limit for any variable constant within tree-holes, and size and shape together explained 20% (*P* = 0.016), that is, 38% of the tree-hole effect (Fig. 2D, Suppl. Table 3 and Suppl. Fig. 6). When the RDA was conducted considering only the first three PCoA axes, these values were 90% and 46% (*P* = 0.007). Time-varying variables explained only 3% of the compositional variance not shared with morphology (Suppl. Note 1). This is consistent with earlier work showing that the number of bacterial taxa in tree-holes increases with volume, as in oceanic islands [43]. A purely descriptive study might have concluded that tree-hole morphology imposes a selective pressure differentiating these communities, and discounted any role for stochastic processes.

**Figure 3.**
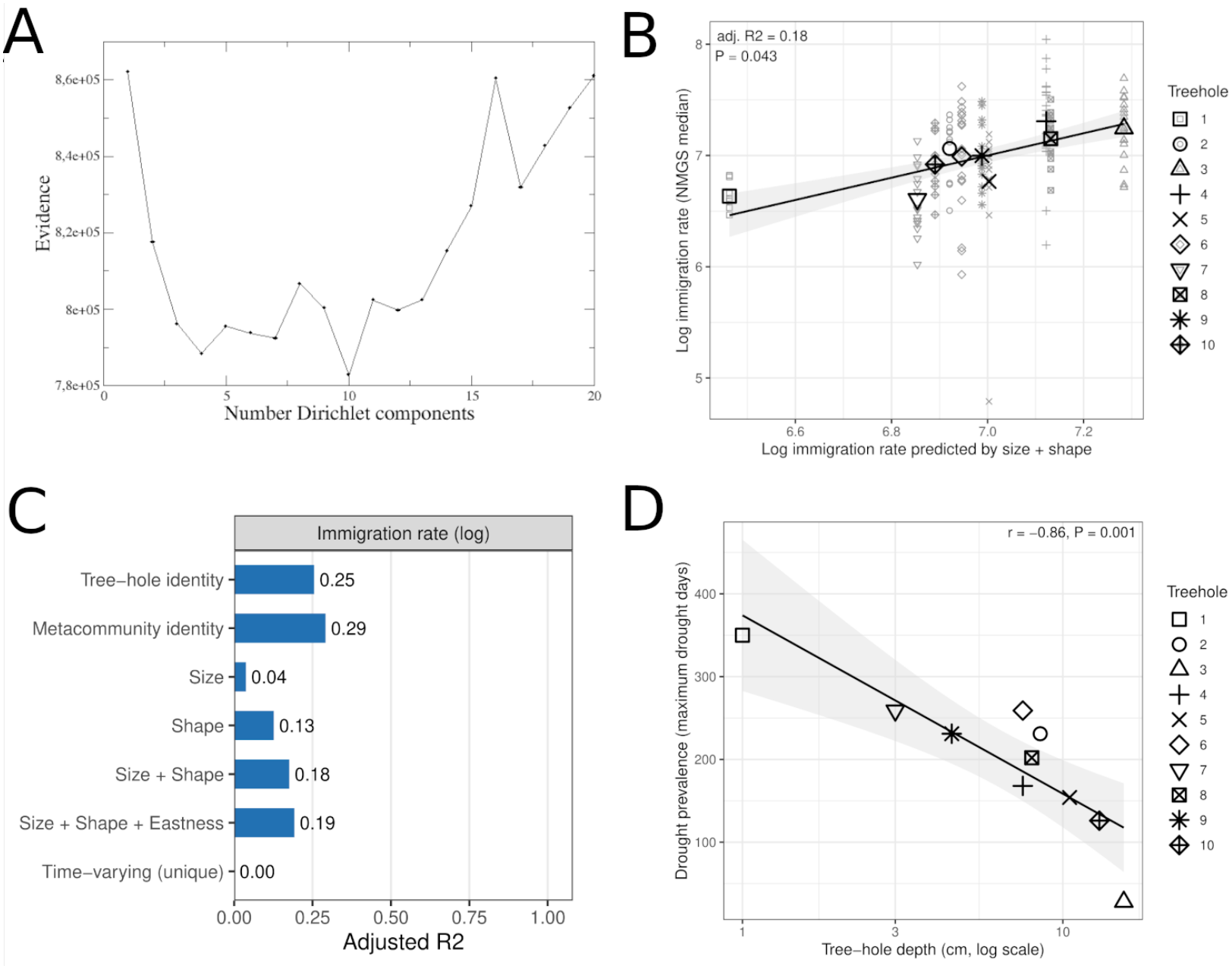
Neutral model parameters and tree-hole morphology. (A) Model evidence of the Dirichlet mixture as a function of the number of components supported the existence of 10 compositional clusters, each interpreted as the set of local communities immigrating from a distinct metacommunity (M1–M10). (B) Log immigration rate inferred for each local community against the value predicted by tree-hole size and shape, the morphological model specified a priori. Small symbols represent samples and large symbols tree-hole means; the line is the least-squares regression of the observed on the predicted values, with its adjusted *R*^2^ and the *P* -value of a test permuting morphology among the 10 tree-holes (the relationship with size and shape separately is shown in Suppl. Fig. 7). (C) Variance explained (adjusted *R*^2^) by tree-hole and metacommunity identity, by tree-hole morphology and by the time-varying variables (unique fraction, given morphology), for the log immigration rate and for the immigration and speciation rates analysed jointly (RDA of both standardised log rates, whose *R*^2^ is the mean of the *R*^2^ of the two rates). (D) Drought prevalence of each tree-hole (maximum number of days that it remained dry during the survey) as a function of its depth; the line is a least-squares fit with 95% confidence band, and *r* is the Pearson correlation with log depth (*n* = 10).

An established strategy to study stochastic processes is Hubbell’s neutral theory [44], in which local communities assemble stochastically through immigration from a shared species pool, the metacommunity, in which new species arise at a given (speciation) rate, and individuals of all species have equal birth and death rates. Species abundance distributions generated in this way serve as a null model against which deterministic hypotheses, such as environmental filtering or ecological interactions, can be tested. We fitted our data to a multi-site neutral model (MSN, see Methods), a computationally tractable implementation of neutral theory for microbial communities [25], assuming that all local communities immigrate from the same metacommunity, each with its own immigration rate.

The MSN allows two nested hypotheses to be tested. The *local* neutral test takes the metacommunity species abundance distribution inferred from the data as given, without constraining its shape, and asks only whether local communities are compatible with neutral sampling from it. The *full* test additionally requires the metacommunity itself to be neutral, i.e. its species abundance distribution must be compatible with neutral dynamics with a single speciation rate (Suppl. Note 2).

We were unable to reject neutrality at the level of local communities (pseudo p-value = 1), whereas the full neutral model was rejected (pseudo p-value < 4 × 10^−4^). A study focusing on neutral assembly might have concluded that this system is adequately explained by a neutral process, with differences between communities explained by immigration rates, as proposed in [16]. However, the rejection of the full model indicated that non-neutral processes operated on the metacommunity, or that the samples do not share a single source pool, which provides a means to reconcile neutral assembly with niche effects.

### Compositional clustering identifies the metacommunity scale

When non-neutral processes operate, *β*-diversity deviates from a null expectation [6, 45], and communities may form sharply separated compositional clusters. We searched for such clusters by fitting a mixture of Dirichlet distributions [26], penalizing the number of components (see Methods), in which each component can be regarded as a distinct source pool from which a subset of local communities is drawn. Throughout this work, we use the term ‘metacommunity’ in this operational sense: for a given set of samples, it is the source species abundance distribution that the MSN infers for that set. Each cluster thus defines one metacommunity (M1–M10, to distinguish them from the tree-holes TH1–TH10), with its own speciation rate and immigration rates specific to each local community, similar in spirit to the niche–neutral hybrid model of Tang and Zhou [46]. Selective structure separating clusters is thereby attributed to the metacommunity level, and the clustering identifies, from the data alone, the scale at which neutral and niche processes should be tested.

Partitioning the dataset into 10 clusters yielded the best model evidence (Fig. 3A). The clusters overlapped considerably with tree-holes, with 7 out of 10 tree-holes having all their samples within a single cluster (Suppl. Table 4). The two largest tree-holes, TH5 and TH10, shared a single cluster, whereas the samples of the smallest ones (TH1, TH4 and TH9) were distributed across several clusters (Fig. 2B and Suppl. Table 1): larger tree-holes showed greater compositional homogeneity over time. Independent neutral tests on the samples of each cluster (excluding M9, which contained a single sample) were unable to reject neutrality for both local and full tests (Suppl. Table 5), indicating that compositional differences were compatible with neutral assembly from distinct metacommunities. This outcome does not follow trivially from the clustering, because the test evaluates the dispersion of local communities around their metacommunity, not merely their compositional similarity. The number of samples per cluster was above 15 in all cases except M4 and M5, and the test has been reported to retain power to reject non-neutral assembly when the number of reads is high [25].

The speciation rates inferred for each metacommunity (Suppl. Table 6) were not associated with any environ-mental variable after correction for multiple testing (strongest association, drought days: Spearman *ρ* = −0.63, *P* = 0.050, corrected *P* = 0.41, see Suppl. Fig. 8). In the neutral model, this parameter combines the per-birth probability that a new type appears in the metacommunity with the size of the metacommunity, and therefore governs the diversity of the source pool, analogously to Fisher’s *α*. Operationally, it can be read as the rate at which taxa new to the metacommunity enter it, for instance through immigration from a broader regional pool (see Suppl. Results for details). We therefore do not interpret it as evidence of speciation, and the case for selection rests primarily on the phylogenetic and functional analyses reported below.

Immigration rates were weakly associated with tree-hole morphology. Size and shape together explained 18% of their variance (*P* = 0.043; Fig. 3B,C), mainly through shape: immigration rates were higher in tree-holes that were deep relative to their surface (Suppl. Fig. 7). This association was not significant once the selection among alternative morphological models was taken into account (*P*_sel_ = 0.17), and could not be separated from metacommunity identity (Suppl. Note 2). No time-varying variable was associated with immigration rates within tree-holes. In contrast with Rivett *et al*. [16], who related immigration rates to tree-hole surface area, surface area alone did not explain them (*R*_adj_^2^ < 0), and the tree-holes with higher immigration rates were those with a smaller surface relative to their depth. Tree-holes with this shape also dried out less often: drought prevalence (maximum number of drought days during the survey) decreased with tree-hole depth (*r* = −0.86, *P* = 0.001; Fig. 3D and Suppl. Fig. 9), the expected proxy for water retention, since the volume of water per unit of evaporating surface is proportional to depth. Rather than accessibility to immigrants, the inferred immigration rate may therefore partly reflect the capacity of tree-holes to retain water, which can act as an environmental filter on establishment.

### Selection and dispersal limitation act between metacommunities, stochastic processes within them

Species abundance distributions compatible with neutral processes can arise in simulations that include strong selection [47], so confirming selection requires exploring phylogenetic relatedness and traits. We used the test proposed by Dini-Andreote *et al*. [10], based on the *β* nearest taxon index (*βNTI*; see Methods): *βNTI* < 2 indicates homogeneous selection and *βNTI >* −2 variable selection. Otherwise, the Raup–Crick index distinguishes homogenizing dispersal (negative values) and dispersal limitation (positive values), and differences are attributed to drift when neither index is significant.

The prior identification of metacommunities facilitated the interpretation of these results. In the heatmap shown in Fig. 4A, each cell represents a pairwise comparison of communities, with communities belonging to the same metacommunity enclosed in black boxes along the diagonal and comparisons between metacommunities in off-diagonal cells. *βNTI* is shown when significant, indicating that selection was the dominant process; otherwise, the Raup–Crick index is shown.

**Figure 4.**
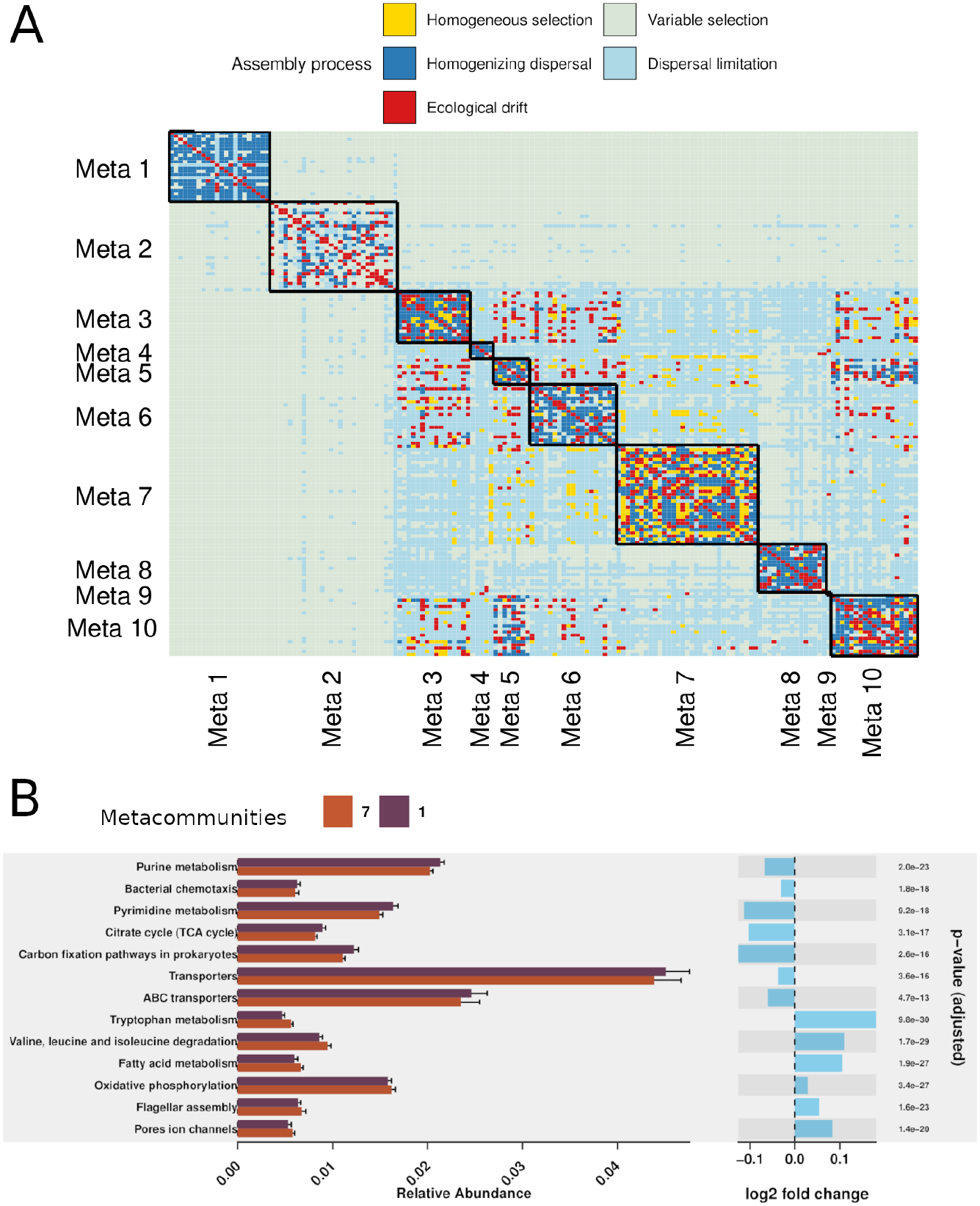
Evidence of selection and dispersal. (A) All-against-all comparison of communities following the statistics proposed by Dini-Andreote *et al*. [10]. Each colour represents evidence of selection, dispersal or drift as indicated (details in the main text). (B) Comparison of the relative abundances of KEGG pathways inferred with PICRUSt for M1 and M7, showing the pathways overrepresented in each metacommunity (more examples in Suppl. Materials).

The strongest signal indicated variable selection between M1 and M2, and between each of them and the remaining metacommunities. These metacommunities contained the three largest tree-holes: TH3 (M1), and TH5 and TH10 (M2) (Suppl. Table 1), which differed in shape: TH3 was deep relative to its surface, whereas TH5 and TH10 were wide relative to their depth (Suppl. Fig. 3), which may explain their different selective pressures. Beyond variable selection, dispersal was homogenizing within metacommunities and limited between them, consistent with stochastic assembly within metacommunities and with physical barriers differentiating them. Within metacommunities, we also observed homogeneous selection, to a lesser extent.

We then investigated whether variable selection translated into differences in gene repertoires by predicting metagenomes with PICRUSt [35], focusing on 25 KEGG pathways previously reported as ecological signatures of tree-hole communities [17]. Metacommunities differentiated by variable selection tended to show distinct sets of pathways (e.g. M1 and M7, Fig. 4B, or M1 and M2 Suppl. Fig. 10), whereas those differentiated only by dispersal limitation yielded fewer differentially abundant pathways, with smaller effect sizes (e.g. M3 and M7, Suppl. Fig. 10). Since dispersal limitation cannot be unambiguously classified as deterministic or stochastic [48], these differences need not reflect adaptation, and may arise from restricted dispersal or priority effects.

### Co-occurrence networks recapitulate metacommunity structure

If metacommunities are the appropriate unit for testing neutrality, they should also be the appropriate unit for network inference, because taxa associated with specific environments tend to co-occur, confounding interactions with shared preferences [49, 50]. Ecological interactions act as a selective force, but how they affect microbial composition and, in turn, metrics intended to detect selection such as *βNTI* remains understudied. Rather than using methods that attempt to control for environmental factors [51, 52], we adopted the conservative approach of inferring a separate SparCC network [39] for each cluster [53, 54] (see Methods). If associations largely reflected shared environmental preferences, clusters with higher compositional similarity should yield more similar networks. Indeed, grouping clusters by the links shared by their networks reproduced the grouping obtained from *β*-diversity, with M1 and M2 separating from all others (Fig. 5A). This correspondence emerged only at permissive thresholds yielding hundreds of thousands of links, unlikely to reflect ecological interactions (see Suppl. Results for further details).

**Figure 5.**
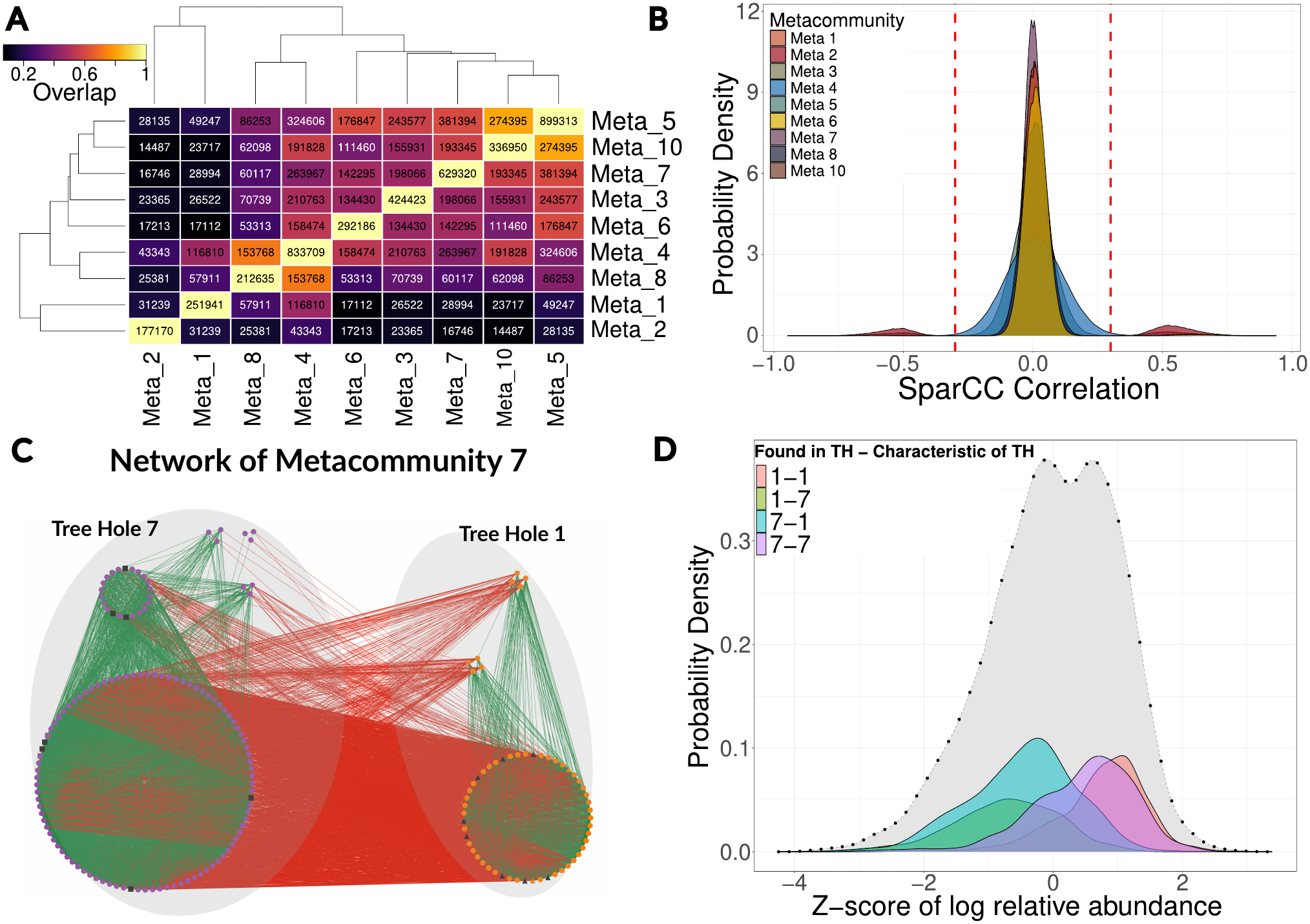
Co-occurrence networks recapitulate metacommunity structure. (A) Szymkiewicz-Simpson similarity (overlap) between SparCC networks inferred for each compositional cluster (metacommunities M1– M10), with the number of shared links indicated in each cell. M9 was excluded from the analysis as it had only one sample. (B) Probability densities of SparCC scores, showing significantly high correlations only for two metacommunities (M2 and M7), with the effect-size threshold indicated as a dashed line. (C) Co-occurrence network of M7 for taxa selected via IndVal and Rank-Abundance. Groups indicate functionInk guilds; only those with more than 3 ASVs are shown. Edge colors: green (positive), red (negative). Node colors denote rank-abundance selection (orange: TH1, purple: TH7, black: no preference). Node shapes denote IndVal selection (triangles: TH1, squares: TH7, circles: unselected). (D) Standardized log-relative abundance distributions (*z*-score) for M7 and IndVal-selected taxa on each TH (*n* = 12 for TH1; *n* = 19 for TH7).

The same confounding was apparent within a single metacommunity. The network of M7, which comprised TH1 and TH7 and showed the strongest evidence of homogeneous selection (Fig. 4A), included strong correlations (Fig. 5B) organised in two groups, with positive associations within and negative associations between them (Fig. 5C). We hypothesized that, although both tree-holes were compositionally similar, these groups reflected different environmental conditions in each tree-hole. To explore this hypothesis, we searched for taxa characteristic of each tree-hole (see Methods), finding that these groups indeed corresponded to taxa associated to distinct tree-holes. Although most taxa were present in both, they had markedly different abundances in each tree-hole (Fig. 5C,D). Therefore, the most economical hypothesis is that each tree-hole host different environmental conditions, and should not be interpreted as interactions. Only a small subnetwork, dominated by positive associations between ASVs from different classes, was conserved in networks inferred separately for each tree-hole (Suppl. Fig. 11); these associations are the most plausible candidates for interactions, although a shared, unmeasured micro-environmental preference cannot be excluded.

## Discussion

Deterministic and stochastic processes, often presented as a dichotomy, are compatible in this system once the scale at which each operates is identified. Selection and dispersal limitation acted between metacommunities, whereas stochastic processes prevailed within them (Fig. 4A), and local communities were statistically indistinguishable from neutral assembly from their respective metacommunities. In microbial systems, the local communities sharing a metacommunity can rarely be delimited from observed movements of individuals, and unsupervised compositional clustering provided this delimitation from the data. The importance of stochastic processes is still debated [55], partly because applying null and neutral models to hundreds of species was computationally prohibitive until recently. We showed that a comprehensive picture of the framework proposed by Vellend can nevertheless be obtained with existing tools, although limitations remain (see also [56]).

Restricting the analysis to specific tests would have led to partial conclusions. A neutral test alone would have suggested neutral assembly, as reported for this dataset [16] and for tree-holes from a different woodland (Wytham Woods, Oxford, UK) [14], with differences between tree-holes explained by the influence of surface area on immigration [16]. Conversely, a focus on compositional and functional differences would have suggested that selection dominates, as in Pascual-García & Bell [17]. These studies are not erroneous: each captured part of a multi-scale picture, and their disagreement is resolved once the level at which selection and neutrality operate is made explicit. Our analysis also refines their interpretation: immigration rates were not associated with surface area but tended to increase with depth relative to surface, a shape associated with increasing water retention and, hence, fewer drought days. Although weak and confounded with metacommunity identity, this association suggests that apparent differences in immigration partly reflect a niche effect on establishment rather than differences in propagule supply.

It is also necessary to combine methods because they may differ in their sensitivity with regard to sampling efforts. For instance, restricting Stegen’s test to a single metacommunity could have erased the evidence for variable selection, not only because fewer environmental conditions would be sampled, but because its null model depends on the taxa present, and fewer samples reduce the phylogenetic diversity available. Similar concerns apply to other metrics that depend on the number of samples, such as the determination of phylogenetic core groups [57]. The failure to reject neutrality within clusters may also depend on the number of samples and on sequencing depth, and is not proof of neutrality. Unsupervised clustering, which is computationally inexpensive and, with a Dirichlet mixture, appropriate for compositional data, mitigates these issues: it delimits sets of communities likely subject to similar selective pressures. The number of clusters may help to reveal the intrinsic scale of the system, suggesting whether further sampling is needed if, for example, a single cluster is found, or how strong and homogeneous selection is (if few clusters with many samples each are found).

Exploring the relationship between clusters and scale is particularly useful if spatial, temporal or host metadata is available. In this regard, compositional clustering may be particularly important in complex environments, such as when environmental gradients or multiple scales exist. The balance between deterministic and stochastic assembly often shifts along gradients, with selection prevailing towards the extremes and stochasticity peaking at intermediate conditions, both in space, as along the mesohaline-to-hypersaline salinity range [58], and in time, as in soil communities exposed to experimental dry–wet cycles [59]. The inferred prevalence of each process may also depend on the temporal resolution of sampling [10], particularly in variable environments such as soils [60], freshwater [61] or seasonal marine systems [62]. The closest parallel to our results comes from sinking marine particles, for which selection at depth was detected only at the metacommunity level, with local communities consistent with neutral assembly [63], whereas communities assembling on synthetic particles under controlled conditions rejected neutrality throughout a reproducible succession [64]. In the latter example, compositional clustering was necessary to distinguish different selection stages. Host-associated communities such as the gut are also subjected to nutrient pulses. Traditionally regarded as deterministic [47, 65], increasingly show a combination of niche and neutral effects [66, 67, 68], with shifts towards determinism following changes in diet or host age [69, 70, 71].

The same scale dependence affects the inference of ecological interactions: co-occurrence networks recapitulated metacommunity structure and, within a metacommunity, tree-hole identity (Fig. 5). Since even methods that control for confounding factors [51] cannot establish that associations correspond to interactions, a conservative strategy is to infer separate networks for each compositional cluster, or metadata-defined group [53, 72]. Yet determining that associations inferred are true interactions require independent evidence coming from (e.g.) metabolic models [73, 74, 75], or experiments [76].

Another challenge is demonstrating the existence of selective forces because, most analyses, including ours, rely on community structure. *β*NTI and predicted metagenomes supported selection at the metacommunity level, but predicted pathways do not demonstrate that specific traits are selected, and operate at the community level, which makes it difficult to disentangle the contribution of individual taxa [77]. This contribution may become more complex in the presence of strong interactions. For instance, if cross-feeding is pervasive in natural environments [78], it may lead to the formation of complementary consortia [79]. Complementary (instead of redundant) traits observed from coexisting members that may be phylogenetically divergent may lead to a statistically weak signal for traits involved in these interactions. Recent macroecological approaches, in which stochastic Lotka–Volterra models with community-level interaction patterns reproduce the correlations observed in real data [80, 81], may offer a mechanistic basis for null models including interactions, and a connection with modern coexistence theory [82].

In summary, microbial ecology already has a variety of methods to deliver comprehensive analysis exploring deterministic and stochastic forces acting in natural microbiomes, but more work is needed to integrate these tools and to fill some gaps such as the influence of ecological interactions. Our work points to an accessible approach: the identification of distinct compositional clusters of communities determines an intrinsic scale of the system, upon which a range of analyses such as those presented can be applied to resolve the determinism–stochasticity dichotomy.

## Supporting information

Supplementary Methods and Results

## Software and Data Availability

The software and data required to reproduce the analysis can be found in the repository https://github.com/apascualgarcia/niche_vs_neutral.

## Acknowledgments

We thank Damian Rivett for sharing the raw sequencing data and metadata to perform the analysis. We thank Chris Quince for advice in the use of the multi-site neutral model software.

## Author contributions

Conceptualization: APG. Data curation: JR, APG. Formal analysis: APG. Investigation: all authors. Methodology: JCC, JR, APG. Visualisation: JCC, JR, DA, APG. Software: JCC, JR, DA, APG. Supervision: TB, APG. Validation: JR, APG. Writing (original draft) APG. Writing (review & editing): JR, TB, APG. All authors approved the final version.

