## Supplementary Methods and Results for "Identifying the metacommunity scale reconciles deterministic and stochastic assembly in a natural mainland–island microbial system"

November 15, 2026

(1) Department of Life Sciences, Imperial College London, Silwood Park Campus, Ascot, UK; (2) Centre for Cancer Biomarkers and Biotherapeutics, Barts Cancer Institute, Queen Mary University of London, London, UK; (3) Centre for Advanced Studies of Blanes (CSIC), Blanes, Spain; (4) Spanish National Centre for Biotechnology (CSIC), Madrid, Spain;

#### Supplementary Methods

**Bioinformatics pipeline.** Raw sequencing data were retrieved from the authors of Ref. [1]. We demultiplexed the raw paired-end reads with idemp (`https://github.com/yhwu/idemp`) using the index reads and sample barcodes, and processed them with DADA2 v1.32.0 [2]. Reads containing ambiguous bases were first removed. Primers 515F (5'-GTGYCAGCMGCCGCGGTAA-3') [3] and 806RB (5'-GGACTACNVGGGTWTCTAAT-3') [4] were then trimmed from both reads with cutadapt v4.6 [5], and reads shorter than 50 bp after trimming were discarded. Forward and reverse reads were truncated at 120 and 220 bp, respectively, and reads with more than two expected errors or matching the PhiX genome were removed. Error rates were learned separately for forward and reverse reads from $10^8$ bases of randomly selected reads, and amplicon sequence variants (ASVs) were inferred with sample pooling. After merging the read pairs, chimeric sequences were removed with the consensus method (98.3% of the reads were retained). Taxonomy was assigned with the naive Bayesian classifier [6] against the SILVA v138.1 database [7]. ASVs classified as chloroplasts or mitochondria were removed, as well as samples with 5000 reads or fewer, ASVs with fewer than 100 reads in total or present in fewer than 10 samples, and the negative controls, leaving 3537 ASVs in 164 samples.

**Test of neutrality.** We tested whether the communities were compatible with neutral assembly using the multi-site neutral model of Harris *et al.* [8]. In brief, the method follows a Bayesian approach that independently fits the matrix $\mathbf{X}$ describing the abundances of ASVs in the samples belonging to a given set (either all samples or the samples of one compositional cluster, each set defining one metacommunity) to a Hierarchical Dirichlet Process with speciation parameter $\theta$, metacommunity's distribution $\bar{\beta}$ and immigration rates $I_i$ $(i = 1, \ldots, M)$, with $M$ the number of samples in the set. Posterior samples of parameters, labelled with the index $k$, i.e. $\bar{\beta}^k, \theta^k, I_1^k, \ldots, I_M^k$, are then obtained. Next, synthetic matrices $\{\mathbf{X}_0^k\}$ with the same number of samples and abundances as the set under study were sampled from each set of parameters $k$, simulating a Hierarchical Dirichlet Process (HDP). Finally, the metacommunity distribution parameters of each synthetic matrix are inferred $(\bar{\beta_0}^k)$. The method allows for the generation of neutral local communities, while the metacommunity may or may not be neutrally assembled. In the local neutral test, the metacommunity distribution $\bar{\beta}$ is left unconstrained, so that only the sampling of local communities from it is required to be neutral. In the full neutral test, $\bar{\beta}$ is additionally required to be consistent with a neutral metacommunity, i.e. with a draw from a Dirichlet process with concentration (speciation) parameter $\theta$. To test for neutrality, 50000 realizations of posterior samples were generated with 25000 being discarded as burn-in, and then 2500 selected among the last 25000 in intervals of 10 steps. The test considers that the observed data are neutral if the proportion of the log-likelihoods $L(\mathbf{X}|\bar{\beta}^k, I_1^k, \ldots, I_M^k)$ estimated from the observed data exceeds the corresponding log-likelihood estimated with synthetic data $L(\mathbf{X}_0^k|\bar{\beta_0}^k, I_1^k, \ldots, I_M^k)$ by some amount. This proportion is an empirical (pseudo) p-value and we considered neutrality rejected if $p < 0.001$. Because the power of neutral tests depends on the number of samples, tests were restricted to clusters with more than ten samples (cluster M9, with a single sample, was excluded); the multi-site model has been reported to remain discriminating in this regime when the number of reads is large [8]. Computations were performed using the code provided in Ref. [8].

**Inference of metacommunities.** We performed an unsupervised classification of samples using a Bayesian approach proposed by Holmes *et al.* [9]. The approach considers the matrix of observations $\mathbf{X}$ whose entries $X_{ij}$ describe the abundance of ASV $j \in \{1, \ldots, S\}$ in community $i \in \{1, \ldots, M\}$, and it assumes that each community is drawn from a multinomial distribution with a vector of $S$ parameters $\bar{p}_i$. In a Bayesian framework, a prior distribution must be chosen for these parameters, and the Dirichlet distribution $\mathrm{Dir}(\bar{p}_i|\bar{\alpha})$ is a natural choice, since it is the conjugate prior of the multinomial distribution, so that the posterior distribution is also a Dirichlet. The procedure then fits the data to infer the parameters $\bar{p}_i$ and $\bar{\alpha}$. The vector of $S$ parameters $\bar{\alpha}$ may be common to all samples, or there may be $k$ subsets of samples, each with a different vector $\bar{\alpha}_k$. In the latter scenario each vector of $\bar{\alpha}_k$ parameters can be interpreted as a different 'metacommunity' $k$ from which a subset of samples was drawn, a possibility that we exploited in our analysis. To perform the fit, the model considers a linear combination of Dirichlet distributions $\mathrm{Dir}(\bar{p}_i|\bar{\alpha}_k)$, providing a flexible framework to fit subsets of communities to distributions with different parameters. To prevent overfitting, models with more components were penalized by selecting the number of components that maximizes the model evidence [9], which determined the number of clusters. Each cluster (the subset of communities assigned to the same Dirichlet component) defines a metacommunity, which we labelled M1–M10. Performing an independent neutral test on each cluster is, in this framework, equivalent to positing a distinct metacommunity per cluster; selective structure separating clusters is thereby attributed to the metacommunity level rather than to local assembly. The fit was performed with the software provided in the original article [9].

**Environmental drivers of composition and of neutral-model parameters.** ASV counts were not rarefied. Size closely follows log volume and shape the log depth-to-surface ratio; the exact relationship is derived in Suppl. Section  and shown in Suppl. Fig. 4. In the max-statistic correction [10], the maximum adjusted $R^2$ over all combinations of one to three morphological variables was recorded in each permutation, while keeping the composition and the tree-hole membership of each sample. Time-varying variables were tested with 999 permutations of samples within tree-holes, and the variance was partitioned between morphology and time-varying variables following [11]. For the immigration rates, we used the median of the posterior distribution (one value per sample), and the analysis was complemented by a linear mixed model with a random intercept per tree-hole [12]. Multiple-testing correction of the correlations with the speciation rates followed Benjamini and Hochberg [13].

**Raup–Crick dissimilarity.** When the $\beta NTI$ comparison yielded a non-significant value, we computed the Raup–Crick dissimilarity. For each pair of sampling units, we computed the observed number of shared species ($s_{\mathrm{obs}}$) and compared it to a null distribution of overlap values ($s_{\mathrm{null}}$) generated by randomizing species occurrences while maintaining fixed row and column sums (i.e., fixed community richness and species incidence). Specifically, we computed the standardized probability that randomized communities produce an overlap smaller than or equal to the observed overlap:

$$p_{ij} = \frac{\#\{s_{\mathrm{null}} \leq s_{\mathrm{obs}}\} + 0.5\,\#\{s_{\mathrm{null}} = s_{\mathrm{obs}}\}}{N_{\mathrm{sim}}}$$

where $N_{\mathrm{sim}}$ is the number of null replicates and ties were handled by assigning half weight to $s_{\mathrm{null}} = s_{\mathrm{obs}}$. The Raup–Crick dissimilarity was then scaled as $RC_{ij} = 2p_{ij} - 1$. Values of $RC_{ij} < 0$ are interpreted as indicators of homogenizing dispersal (more similar than expected by chance) whereas $RC_{ij} > 0$ represent the existence of dispersal barriers, with values near 0 indicating drift. Computations were conducted with the R package iCAMP [14] using the function bNTIn.p with 1000 randomizations.

**Inference of associations.** SparCC significance was assessed through 1000 bootstrap permutations; only correlations achieving a pseudo $p$-value $< 0.001$ were considered statistically significant, and correlations with an absolute SparCC coefficient $|r| < 0.3$ were discarded. The Szymkiewicz-Simpson overlap coefficient ($O_{AB}$) measures the proportion of shared edges relative to the size of the smaller network. In M7, Indicator Value Analysis (IndVal) was performed with the `indval` function from the `labdsv` R package [15] with the `duleg` parameter set to `TRUE`, restricting the search to strict indicator ASVs, and significance was assessed with 1000 Monte Carlo permutations, retaining ASVs with a pseudo $p$-value $\leq 0.05$. We then relaxed the IndVal criteria by also including ASVs with $|\Delta\text{Abundance}| > 0.05$, or $|\Delta\text{Rank}| > 1000$. Although the latter criteria are permissive, the additional signal outweighed the noise introduced by rare taxa with small differences in relative abundance (Suppl. Fig. 1). To facilitate the visual representation and structural interpretation of the inferred networks, we identified ASVs sharing approximately the same neighbours with the same types of links (guilds) with `functionInk` [16]; the optimal partition of the network into guilds was identified as the one maximizing the total partition density.

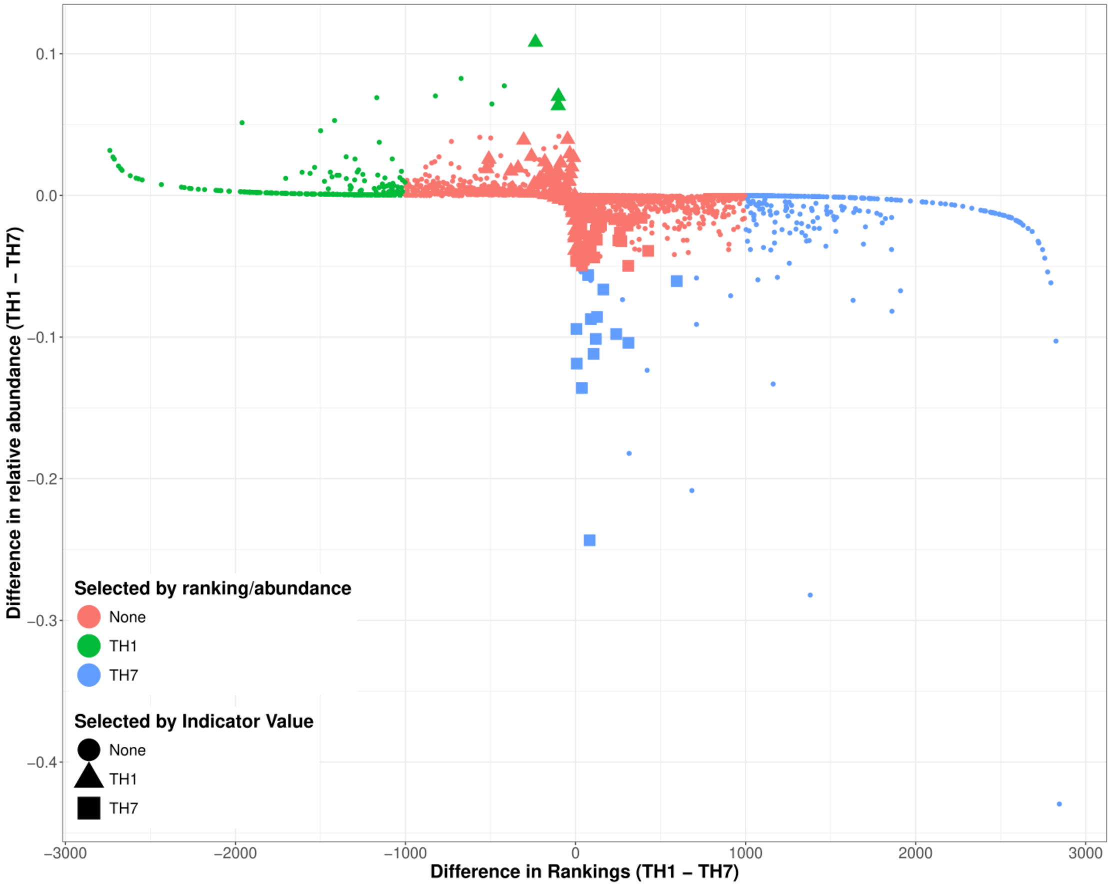


Figure 1: **Compositional difference between TH1 and TH7 within Metacommunity 7**. Relationship between the taxonomic rank shifts and differences in qumulative relative abundance between the two tree holes. Points are color-coded to highlight taxa that exceeded the filtering thresholds ($|\Delta\text{Rank}| > 1000$ or $|\Delta\text{Abundance}| > 0.05$), with distinct colors representing alignment toward one specific TH. Point shapes indicate taxa independently identified as characteristic of the THs via Indicator Value (IndVal) analysis. These taxa are displayed with a bigger size to enhance visual clarity.

**Distance.** The standard Aitchison distance (CLR with a pseudocount of one) was preferred to the robust Aitchison distance implemented in `vegan` [17]. The robust version computes the CLR on non-zero counts only and replaces the resulting matrix by a low-rank reconstruction (rank 3 by default), so that its PCoA has as many axes as the chosen rank. Since zeros are ignored, it discards the information on which ASVs are present, which in this dataset carried the main compositional gradient (Suppl. Section ).

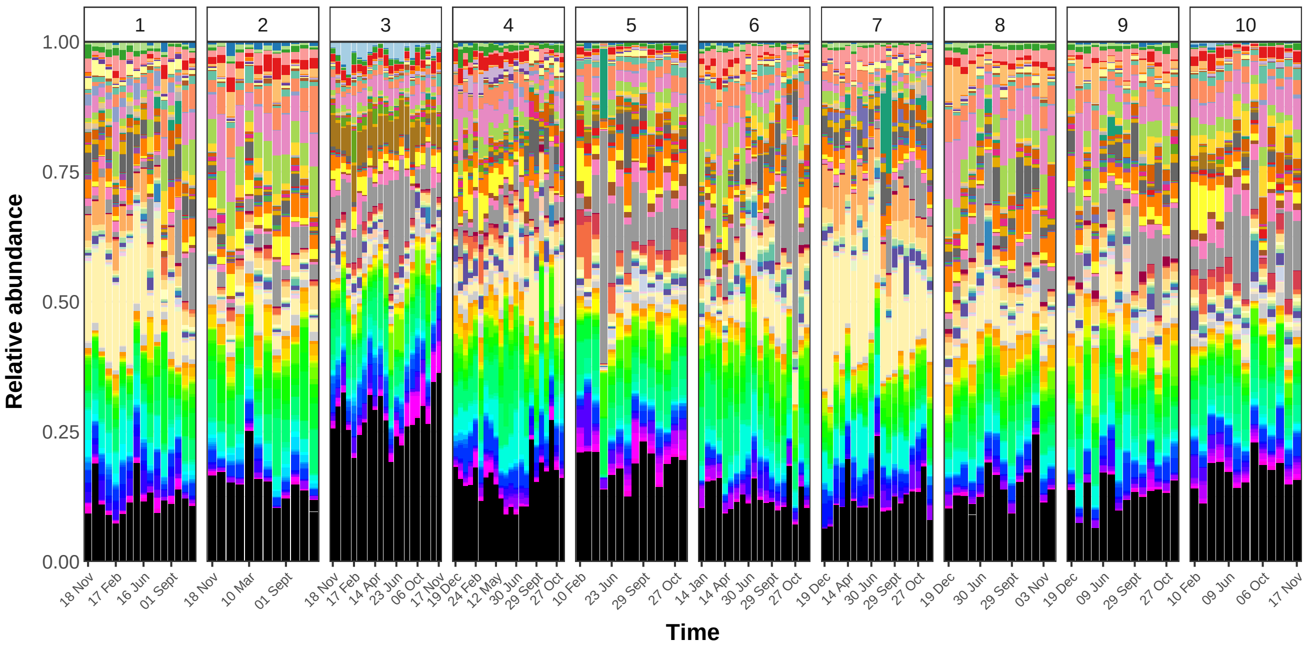


Figure 2: **Tree-holes composition at the Genus level.** Bar-plots representing ASVs composition for each tree-hole (boxes) ordered by sampling time. Genera representing $> 1\%$ relative abundances are highlighted.

**Permutation tests.** Morphological variables are constant within each tree-hole, so a permutation of samples treats 147 samples as independent replicates of 10 morphological values and overstates significance. The test used here permuted the rows of the table of tree-holes (one row per tree-hole) and assigned to each sample the morphology of the tree-hole that occupied its position; composition and tree-hole membership were not modified, and the correlations among morphological variables were preserved. Because the number of samples per tree-hole varied (8 to 21), this test was implemented explicitly rather than with the block permutations of the `permute` package [18]. Time-varying variables were tested by permuting samples within tree-holes; this test does not account for the temporal autocorrelation of consecutive samples and may be slightly liberal.

**Model selection.** All combinations of one to three morphological variables (size, shape, log depth, log surface area, log volume, log depth-to-surface ratio, northness and eastness) were fitted, excluding combinations with more than two size-related variables, which are exactly collinear, and pairs other than size and shape, which have the same fit. In each permutation all 30 combinations were refitted, and the corrected $P$-value of a combination was the fraction of permutations in which the maximum adjusted $R^2$ exceeded the observed value [10]. A forward selection [19] was used only as an exploratory screening; it selected size, eastness and shape as the first three variables, in agreement with the exhaustive search.

### Supplementary Note 1. Environmental variables and composition

**Tree-hole morphology.** The physical descriptors of the tree-holes are described in Suppl. Table 1. The four variables indicated were not independent because volume was computed as $V = \text{SA} \times \text{D}/3$, where $SA$ is surface area and $D$ is depth. These four variables therefore lie in a plane, and any model including two of them has the same fit. Within this plane, log depth and log surface area were correlated across tree-holes ($r = 0.72$), and so were log volume and the log depth-to-surface ratio ($r = 0.37$). Since, on a logarithmic scale, $\log(V) = \log(SA) + \log(D) - \log(3)$ and $\log(D/SA) = \log(D) - \log(SA)$, the rotation that defines size and shape (Methods) yields two uncorrelated variables that span the same plane (Suppl. Fig. 3).

Size and shape together are thus the same model as depth and surface area, or as volume and the depth-to-surface ratio, but the effect of each variable can be interpreted separately. Depth is a combination of both dimensions, $z_D = (\text{size} + \text{shape})/\sqrt{2}$, which explains why it had the highest correlation with PCoA1, an axis

| | TH1 | TH2 | TH3 | TH4 | TH5 | TH6 | TH7 | TH8 | TH9 | TH10 |
|---|---|---|---|---|---|---|---|---|---|---|
| volume ($cm^3$) | 25.5 | 247.92 | 284.17 | 92.5 | 304.5 | 170 | 145 | 104 | 45 | 736.7 |
| surface ($cm^2$) | 8.5 | 87.5 | 55 | 37 | 87 | 68 | 28 | 39 | 30 | 170 |
| depth ($cm$) | 1 | 8.5 | 15.5 | 7.5 | 10.5 | 7.5 | 3 | 8 | 4.5 | 13 |
| depth/surface ($cm^{-1}$) | 0.12 | 0.10 | 0.28 | 0.20 | 0.12 | 0.11 | 0.11 | 0.20 | 0.15 | 0.08 |

Table 1: **Tree-holes morphology.** Morphological measurements (depth and surface) and metrics derived from them (volume and depth/surface).

mainly associated with size with a minor contribution of shape, while it was not associated with PCoA3, the axis associated with shape (Suppl. Table 2).

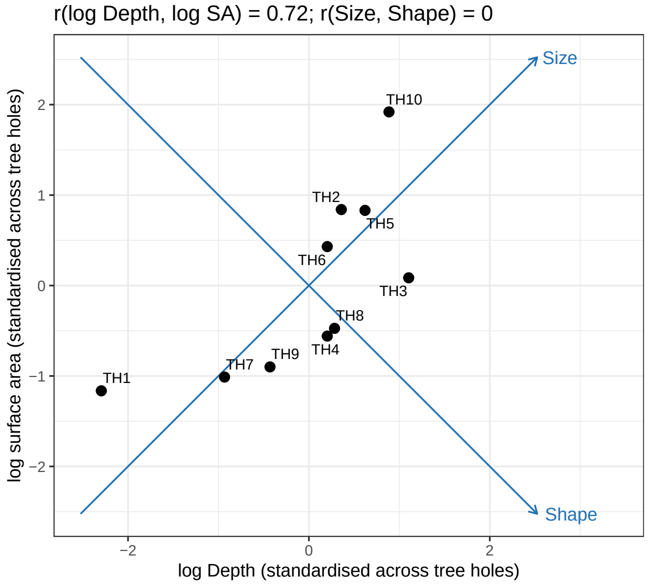


Figure 3: **Definition of tree-hole size and shape.** Each point represents a tree-hole, positioned by its log depth and log surface area, both standardised across the 10 tree-holes. Size and shape are the two diagonals of this plane: size increases with both depth and surface area, and shape increases with depth relative to surface area.

**Size and shape versus the original variables.** Let $\mu_D$, $\sigma_D$ and $\mu_S$, $\sigma_S$ be the mean and standard deviation of log depth ($D$) and log surface area ($SA$) across the 10 tree-holes, so that $z_D = (\log D - \mu_D)/\sigma_D$ and $z_S = (\log \text{SA} - \mu_S)/\sigma_S$. Inverting the definitions of size and shape gives $z_D = (\text{size} + \text{shape})/\sqrt{2}$ and $z_S = (\text{size} - \text{shape})/\sqrt{2}$, and substituting in $\log V = \log D + \log \text{SA} - \log 3$ and $\log(D/\text{SA}) = \log D - \log \text{SA}$ yields

$$\begin{aligned} \log V &= (\mu_D + \mu_S - \log 3) + a\,\text{size} + b\,\text{shape}, \\ \log(D/\text{SA}) &= (\mu_D - \mu_S) + b\,\text{size} + a\,\text{shape}, \end{aligned} \qquad a = \frac{\sigma_D + \sigma_S}{\sqrt{2}}, \quad b = \frac{\sigma_D - \sigma_S}{\sqrt{2}}.$$

With $\sigma_D = 0.81$ and $\sigma_S = 0.62$, $a = 1.01$ and $b = 0.14$ (intercepts 4.71 and $-2.10$, units as in Suppl. Table 1). Log volume is therefore almost a pure function of size, and the log depth-to-surface ratio of shape; the small cross-term $b$ arises because log depth varies slightly more than log surface area across tree-holes. Across tree-holes, size was correlated with log volume ($r = 0.998$) and shape with the log depth-to-surface ratio ($r = 0.95$), whereas the cross-correlations were weak (size with the log depth-to-surface ratio, $r = 0.31$; shape with log volume, $r = 0.05$; Suppl. Fig. 4). The correlation between shape and the log depth-to-surface ratio is lower than that between size and log volume because shape varies less than size (variances $1 - \rho$ and $1 + \rho$, with $\rho = 0.72$ the correlation between log depth and log surface area), so the same cross-term has a larger relative weight.

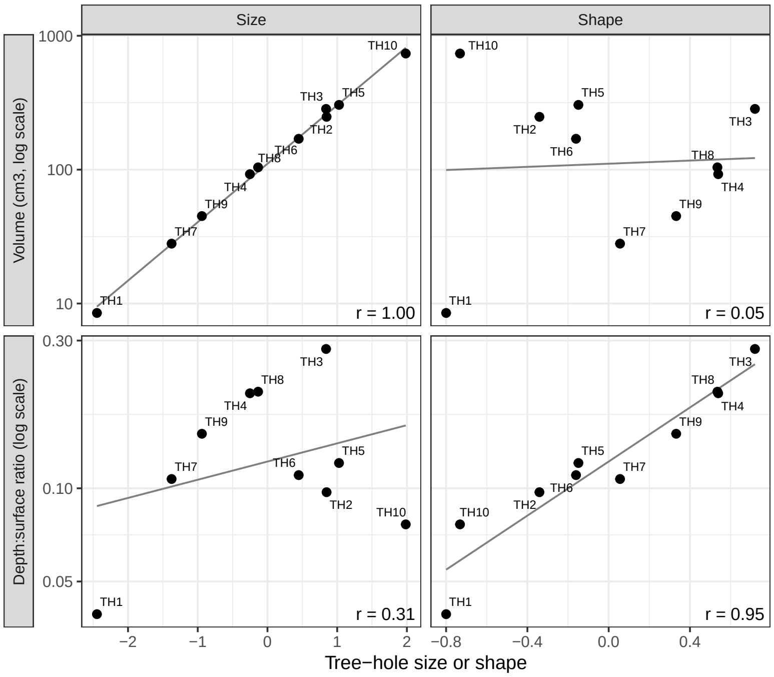


Figure 4: **Size and shape versus the original morphological variables.** Volume (top) and depth-to-surface ratio (bottom), on a logarithmic scale, against tree-hole size (left) and shape (right). Each point is a tree-hole; lines are least-squares fits and $r$ is the Pearson correlation across the 10 tree-holes.

**Composition along the PCoA axes.** The first three PCoA axes explained 21.2%, 12.5% and 8.8% of the compositional variance (Suppl. Fig. 5). The first axis mainly reflected which ASVs were present in each sample: it was almost identical to the first axis of a PCoA of presence–absence (Jaccard) distances ($r = 0.98$), which was itself correlated with log depth ($r = 0.76$). Accordingly, a distance that ignores zero counts (the robust Aitchison distance; Suppl. Section ) did not recover the association of the first axis with depth.

**Morphological models.** Suppl. Table 3 reports the compositional variance explained by tree-hole morphology. Size alone was significant, whereas shape alone was not, because shape was mainly associated with PCoA3, an axis that accounts for a small fraction of the compositional variance. Among the 30 combinations of one to three morphological variables, the highest fit was obtained with size, shape and eastness, but the improvement over size and shape was not significant after correcting for the selection of the best combination.

**Time-varying variables and variation partitioning.** In a partial RDA conditioned on tree-hole identity, with permutations restricted within tree-holes, moisture ($F = 2.06$, $P = 0.001$), the number of drought days ($F = 2.13$, $P = 0.001$) and the maximum air temperature ($F = 1.53$, $P = 0.011$) were significant, whereas the minimum air temperature ($P = 0.055$), rainfall ($P = 0.065$) and light ($P = 0.17$) were not. Each significant variable explained approximately 0.5% of the total compositional variance. In a variation partitioning, size and shape explained 17% of the compositional variance, the three significant time-varying variables 3%, and their shared effect 3% (77% unexplained); with tree-hole identity instead of morphology, the corresponding fractions were 50%, 3% and 3% (44% unexplained). The shared fraction reflects that larger and deeper tree-holes had fewer drought days. A final RDA including size, shape and the three time-varying variables explained 23% of the compositional variance; its first axis was associated with size, the second with shape and the third with the hydric state of the tree-holes (Suppl. Fig. 6).

| | ENVFIT $r^2$ | | $R^2_{\text{adj}}$, single axis | | |
|---|---|---|---|---|---|
| Variable | Axes 1–2 | Axes 1–3 | PCoA1 | PCoA2 | PCoA3 |
| log depth | 0.78 | 0.78 | 0.58 | 0.09 | 0.02 |
| log volume | 0.49 | 0.63 | 0.55 | 0.16 | 0.01 |
| log surface area | 0.39 | 0.55 | 0.33 | 0.20 | 0.17 |
| log depth-to-surface ratio | 0.29 | 0.72 | 0.16 | 0.00 | 0.45 |
| Orientation | 0.04[a] | 0.39 | 0.14[b] | 0.01[b] | 0.39[b] |
| Drought days | 0.21 | 0.21 | 0.18 | 0.00 | 0.01 |
| Moisture | 0.13 | 0.13 | 0.13 | 0.00 | 0.00 |
| Size | – | – | 0.53 | 0.17 | 0.02 |
| Shape | – | – | 0.03 | 0.03 | 0.54 |

[a] $P = 0.052$. [b] Eastness.

Table 2: **Environmental variables associated with the Aitchison PCoA.** Squared correlation ($r^2$) of each variable with the planes of axes 1–2 and 1–3 (ENVFIT; $P = 0.001$ for all variables listed except where indicated), and adjusted $R^2$ of the linear regression of each PCoA axis on each variable. Size and shape were not included in ENVFIT. Temperature, rainfall, light and biomass were not significant (ENVFIT $P \geq 0.10$).

| | CLR (all ASVs) | | | PCoA1–3 | | |
|---|---|---|---|---|---|---|
| Model | $R^2_{\text{adj}}$ | $P$ | $P_{\text{sel}}$ | $R^2_{\text{adj}}$ | $P$ | $P_{\text{sel}}$ |
| Tree-hole identity (upper limit) | 0.53 | – | – | 0.90 | – | – |
| Size | 0.13 | 0.010 | – | 0.32 | 0.007 | – |
| Shape | 0.06 | 0.40 | – | 0.14 | 0.27 | – |
| Size + shape (a priori) | 0.20 | 0.016 | – | 0.46 | 0.007 | – |
| Size + shape + eastness (best) | 0.25 | 0.034 | 0.091 | 0.52 | 0.026 | 0.073 |

Table 3: **Compositional variance explained by tree-hole morphology (RDA).** Adjusted $R^2$ ($R^2_{\text{adj}}$) for the CLR matrix (all ASVs) and for the first three PCoA axes. $P$: test in which the morphology was permuted among the 10 tree-holes (9999 permutations). $P_{\text{sel}}$: $P$-value corrected for the selection of the best of 30 combinations.

### Supplementary Note 2. Neutral-model parameters and environment

**Metacommunities and null model statistics** The number of metecommunities inferred equals the number of tree-holes, with a distribution of tree-holes samples in the metacommunities displayed in Suppl. Table 4. Only three tree-holes have their samples distributed across more than one metacommunity. The number of samples in each metacommunity, and the results of fitting the observed data to a Hierarchical Dirichlet Process are shown in Suppl. Table 5.

**Immigration rates and environmental variables** In a variation partitioning between size and shape and metacommunity identity, morphology alone explained no variance of the log immigration rates ($R^2_{\text{adj}} = -0.01$), the shared fraction was 0.19 and metacommunity identity alone explained 0.11. Since immigration rates are estimated relative to the metacommunity of each sample, and metacommunities were largely nested within tree-holes, the effects of tree-hole shape and of metacommunity identity could not be separated. Within tree-holes, none of the time-varying variables was associated with immigration rates (partial RDA, all $P \geq 0.59$). Among single morphological variables, the log depth-to-surface ratio explained the largest fraction of the variance of the log immigration rate ($R^2_{\text{adj}} = 0.17$, $P = 0.015$, tree-hole permutations, not corrected for selection), followed by shape (0.13, $P = 0.056$) and log depth (0.11, $P = 0.064$); size, log volume and log surface area explained less than 5% (Suppl. Fig. 7).

**Speciation rates.** Suppl. Table 6 reports the strongest correlations between the log speciation rate and the mean of each environmental variable per metacommunity. No correlation was significant after correcting for the 16 variables tested. The correlation with drought days depended partly on metacommunity M9, which comprises a single sample without drought days and had the highest speciation rate (Suppl. Fig. 8); excluding M9, the correlation decreased to $\rho = -0.52$ ($P = 0.16$). The other nine metacommunities had log speciation rates within a narrow range (6.18–6.47). The correlation with log volume, reported in previous versions from a regression

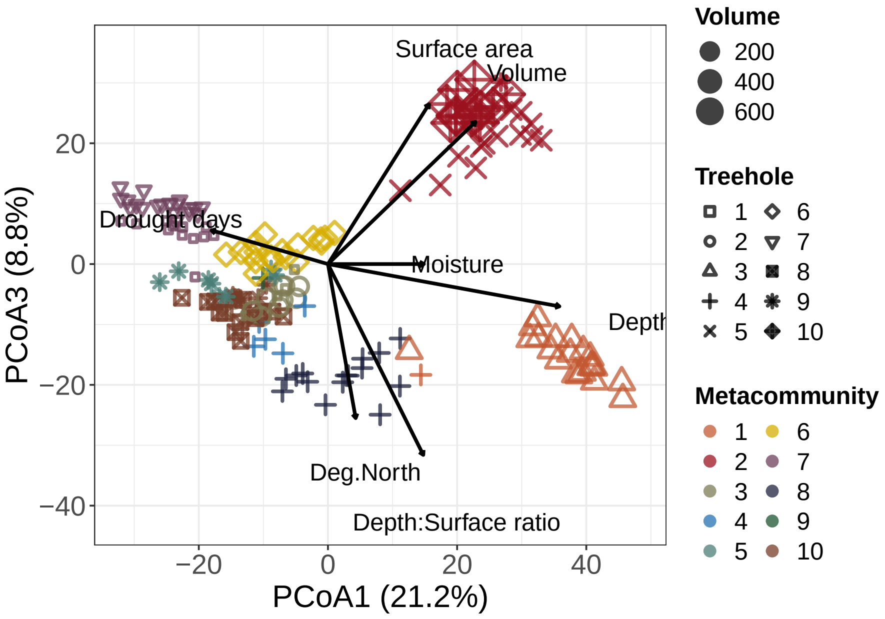


Figure 5: **Aitchison PCoA, axes 1 and 3.** Arrows represent the environmental variables significantly associated with the plane of axes 1–3 (ENVFIT, $P \leq 0.05$). The depth-to-surface ratio and the orientation of the tree-holes were associated with PCoA3.

across samples in which the value of each metacommunity was repeated for all its samples, was not significant at the level of metacommunities ($\rho = 0.35$, $P = 0.32$).

**Drought prevalence and tree-hole morphology.** The number of drought days of a sample is the number of days since its tree-hole was last found holding water (zero if it held water at sampling). We summarised the drought prevalence of each tree-hole as the maximum number of drought days during the survey. Across the 10 tree-holes, drought prevalence decreased with log depth ($r = -0.86$, $P = 0.001$), log volume ($r = -0.75$, $P = 0.012$), size ($r = -0.73$, $P = 0.017$) and the log depth-to-surface ratio ($r = -0.69$, $P = 0.027$), but not significantly with log surface area ($r = -0.49$, $P = 0.15$), shape ($r = -0.49$, $P = 0.15$) or orientation. Depth remained the best predictor when the shallowest tree-hole (TH1, 1 cm deep and dry in 15 of its 16 samplings) was excluded ($r = -0.80$, $P = 0.010$; log depth-to-surface ratio $r = -0.48$, $P = 0.20$), and when drought prevalence was measured as the fraction of samplings in which the tree-hole was dry (log depth $r = -0.87$, log depth-to-surface ratio $r = -0.83$). Depth therefore represents the capacity of tree-holes to retain water better than the depth-to-surface ratio. This is expected from their geometry: since $V = \mathrm{SA} \times D/3$, the volume of water per unit of evaporating surface, $V/\mathrm{SA} = D/3$, is proportional to depth. In terms of size and shape, $z_D = (\mathrm{size}{+}\mathrm{shape})/\sqrt{2}$, so water retention depends on both dimensions of morphology. In contrast, the log depth-to-surface ratio explained more variance of the immigration rates than log depth (see above).

### Supplementary Results

**Composition and environmental variables.** Adding the orientation of the tree-holes did not significantly improve the fit once the selection among alternative combinations of variables was taken into account (Suppl. Table 3). Time-varying variables had a weaker effect than morphology: within tree-holes, moisture and drought days were significant ($P = 0.001$), but together with the maximum air temperature they explained only 3% of the compositional variance not shared with morphology (Suppl. Section ).

**Local and full neutral tests.** The *local* neutral test takes the metacommunity species abundance distribution inferred from the data as given, without constraining its shape, and asks only whether local communities are

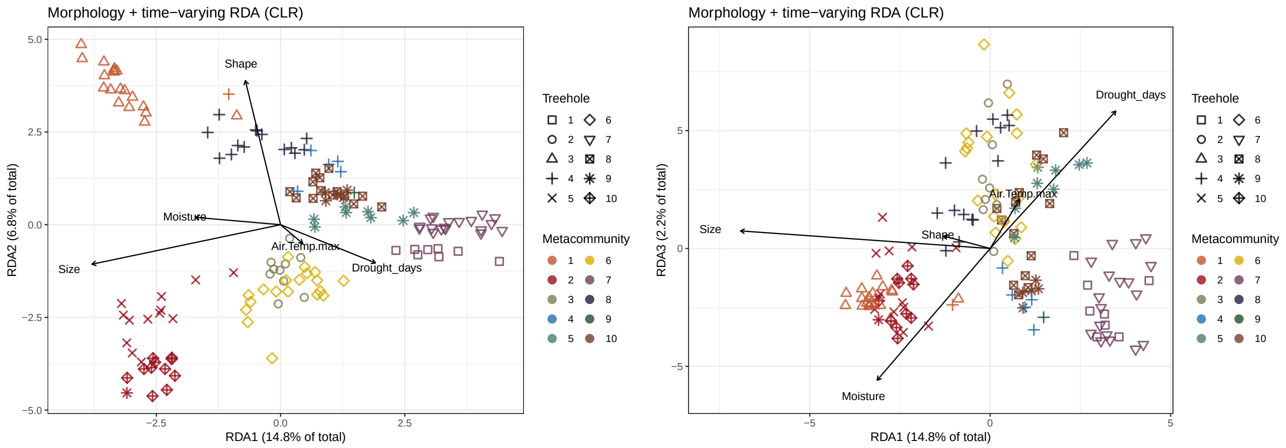


Figure 6: **RDA of composition on tree-hole morphology and time-varying variables.** Axes 1–2 (left) and 1–3 (right). Arrows represent the correlations of the standardised predictors with the constrained axes.

| | M1 | M2 | M3 | M4 | M5 | M6 | M7 | M8 | M9 | M10 |
|---|---|---|---|---|---|---|---|---|---|---|
| TH1 | | | 0.25 | | | | 0.75 | | | |
| TH2 | | | 1.0 | | | | | | | |
| TH3 | 1.0 | | | | | | | | | |
| TH4 | 0.045 | | | 0.23 | | | | 0.68 | 0.045 | |
| TH5 | | 1.0 | | | | | | | | |
| TH6 | | | | | | 1.0 | | | | |
| TH7 | | | | | | | 1.0 | | | |
| TH8 | | | | | | | | | | 1.0 |
| TH9 | | 0.071 | | | 0.57 | | | | | 0.36 |
| TH10 | | 1.0 | | | | | | | | |

Table 4: **Distribution of tree-holes samples in the metacommunities.** Numbers represent the fraction of samples from each tree-hole (TH, rows) that were classified in each metacommunity (M, columns). Empty cells indicate that no samples were classified in the specified combination.

compatible with neutral sampling from it, i.e. whether they differ from the metacommunity and from each other only as expected from drift and finite immigration rates. The *full* neutral test additionally requires the metacommunity itself to be neutral, i.e. that its species abundance distribution is compatible with the one generated by neutral dynamics with a single speciation rate. Rejecting the full model while accepting the local one therefore indicates that local communities sample neutrally from a source pool, but that the relative abundances in this pool were shaped by non-neutral processes, or that the samples do not share a single source pool.

**Rationale of the compositional clustering.** Given the compositional structure described in the main text, it seemed unlikely that differences in immigration rates alone could generate clusters of compositionally dissimilar communities that are sharply separated by an unsupervised method. The rationale is that $\beta$-diversity deviates significantly from a null expectation when non-neutral processes operate [20, 21]. Therefore, if the partition of $\beta$-diversity within and between clusters supports their existence, (possibly different) non-neutral processes act on different subgroups of communities. The Dirichlet mixture used to search for these clusters has a natural interpretation in the context of neutral theory, since each Dirichlet component can be regarded as a distinct source pool from which a subset of local communities is drawn; since adding components always improves the fit, the method penalizes the number of components [9]. In the first neutral test reported in the main text, the set of samples comprised all samples, whereas in the subsequent tests each set comprises the samples assigned to one compositional cluster. Fitting an independent neutral model to each cluster is equivalent to positing a distinct metacommunity per cluster, each with its own speciation rate and with immigration rates specific to each local community.

| Model | Number of samples | Pseudo $p$-val (local/full) | Median immigration rates [Q1, Q3] | Median speciation rates |
|---|---|---|---|---|
| M1 | 22 | 0.71 / 0.73 | [1206.60, 1712.72] | 563.51 |
| M2 | 28 | 0.34 / 1.00 | [860.86, 1167.01] | 611.73 |
| M3 | 16 | 0.70 / 0.99 | [1121.10, 1436.04] | 563.32 |
| M4 | 5 | 0.98 / 0.99 | [1715.52, 2387.33] | 634.45 |
| M5 | 8 | 0.60 / 0.53 | [901.43, 1562.40] | 562.09 |
| M6 | 19 | 0.62 / 1.00 | [875.46, 1577.61] | 623.36 |
| M7 | 31 | 0.13 / 1.00 | [620.35, 869.43] | 484.44 |
| M8 | 15 | 0.74 / 1.00 | [1182.60, 1984.68] | 643.10 |
| M10 | 19 | 0.50 / 1.00 | [999.39, 1400.92] | 609.40 |

Table 5: **Statistics MSN model for each Dirichlet component (metacommunities, M).** For immigration and speciation rates, median were obtained across 25000 sampled parameters. Quantiles of median values across all samples within the same Dirichlet component are shown for immigration rates, whereas there is a single median value for each Dirichlet component.

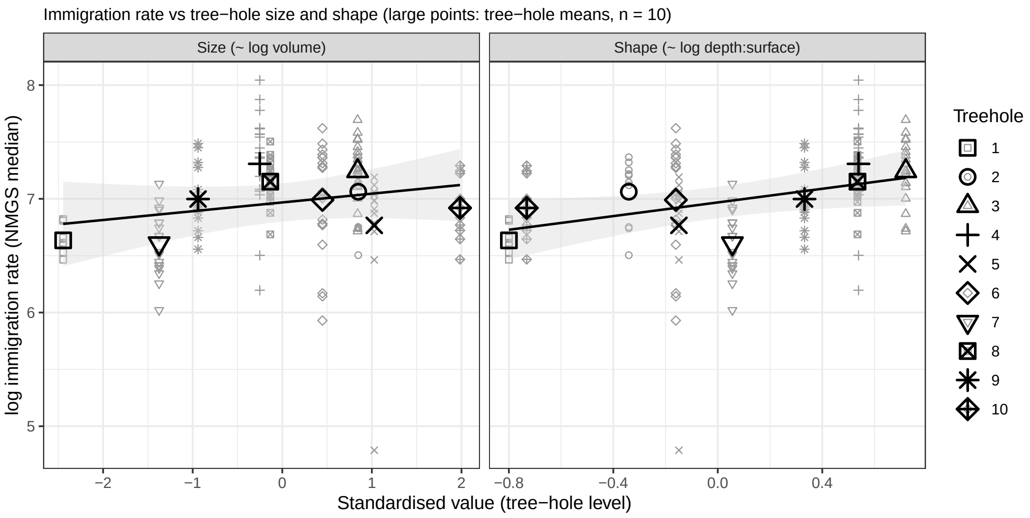


Figure 7: **Immigration rate as a function of tree-hole size and shape.** Small symbols represent samples and large symbols tree-hole means; lines are least-squares fits to the tree-hole means, shown for illustration.

**Neutral tests within clusters.** The failure to reject neutrality within clusters does not follow trivially from the clustering: the multi-site neutral test evaluates the dispersion of local communities around their inferred metacommunity distribution, not merely their compositional similarity, so a cluster that is homogeneous in central tendency can still reject neutrality through over-dispersion or coordinated covariation among taxa. The number of samples per cluster was above 15 in all cases except M4 and M5 (Suppl. Table 5), and the multi-site test has been reported to retain power to reject non-neutral assembly when the number of reads is high [8], with pseudo p-value limits much lower than those obtained here (Suppl. Table 5).

**Interpretation of the speciation rate.** Because each metacommunity has a single speciation rate, we correlated it with the mean of each variable across the samples of the metacommunity ($n = 10$); no correlation remained significant after correction for multiple testing (Suppl. Table 6). This absence of association is consistent with an interpretation in which the inferred speciation rate is unlikely to reflect the emergence of new taxa through evolution. In the neutral model, the parameter $\theta$ is the fundamental biodiversity number, which combines the per-birth probability that a new type appears in the metacommunity with the size of the metacommunity ($\theta = 2J_M\nu$ in Hubbell's formulation [22]); it therefore governs the diversity of the source pool, analogously to Fisher's $\alpha$. Operationally, it can be read as the rate at which taxa that are new to the metacommunity enter it, for instance through immigration from a broader regional pool.

| Variable | $\rho$ | $P$ | $P_{\text{BH}}$ |
|---|---|---|---|
| Drought days | −0.63 | 0.050 | 0.41 |
| Moisture | 0.58 | 0.088 | 0.41 |
| Northness | −0.50 | 0.14 | 0.41 |
| Size | 0.36 | 0.30 | 0.41 |
| log volume | 0.35 | 0.32 | 0.41 |
| Shape | 0.23 | 0.53 | 0.56 |

Table 6: **Spearman correlations between the log speciation rate and the environmental variables** (mean per metacommunity, $n = 10$). $P_{\text{BH}}$: corrected for 16 tests (Benjamini–Hochberg).

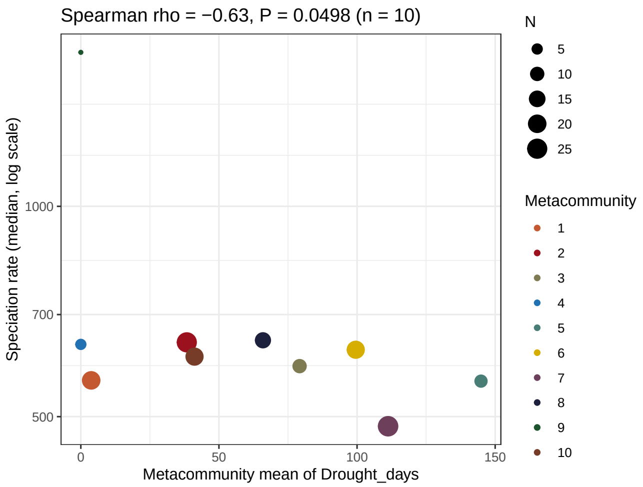


Figure 8: **Speciation rate of each metacommunity as a function of its mean number of drought days.** Symbol size indicates the number of samples per metacommunity; the isolated symbol at the top left corresponds to M9, which comprises a single sample.

**Immigration rates and tree-hole surface area.** The weak association between immigration rates and tree-hole shape reported in the main text should be interpreted with caution for two reasons. First, it was not significant once the selection among alternative combinations of morphological variables was taken into account ($P_{\text{sel}} = 0.17$). Second, it could not be separated from metacommunity identity, which accounted for most of the same variance (Suppl. Section ). The pattern does not support the hypothesis that accessibility to immigrants, determined by tree-hole surface area, shaped these communities [1], since surface area alone did not explain immigration rates and the tree-holes with the highest immigration rates were those with a smaller surface relative to their depth. Among the tree-holes of the metacommunities differentiated by the strongest variable selection (M1 and M2, main-text Fig. 4A), TH3 (M1) was the deepest tree-hole and was deep relative to its surface, whereas TH5 and TH10 (M2) were wide relative to their depth (Suppl. Fig. 3).

**Co-occurrence networks across metacommunities.** Associations inferred from sequencing data should be interpreted with caution, because taxa associated with specific environments tend to co-occur, thereby confounding interactions with shared environmental preferences [23, 24]. We inferred associations within each compositional cluster with SparCC, whose score can be interpreted as a compositionality-corrected Pearson correlation [25], and compared the networks across a range of SparCC thresholds. Grouping clusters by the number of links shared by their networks reproduced the grouping obtained from $\beta$-diversity, with M1 and M2, the metacommunities most strongly differentiated by variable selection (main-text Fig. 4A), separating from all others (main-text Fig. 5A). This correspondence emerged only when the SparCC threshold was so permissive that each network contained hundreds of thousands of links, a number of associations unlikely to reflect ecological interactions. We therefore interpreted them as a consequence of shared environmental conditions or as spurious associations.

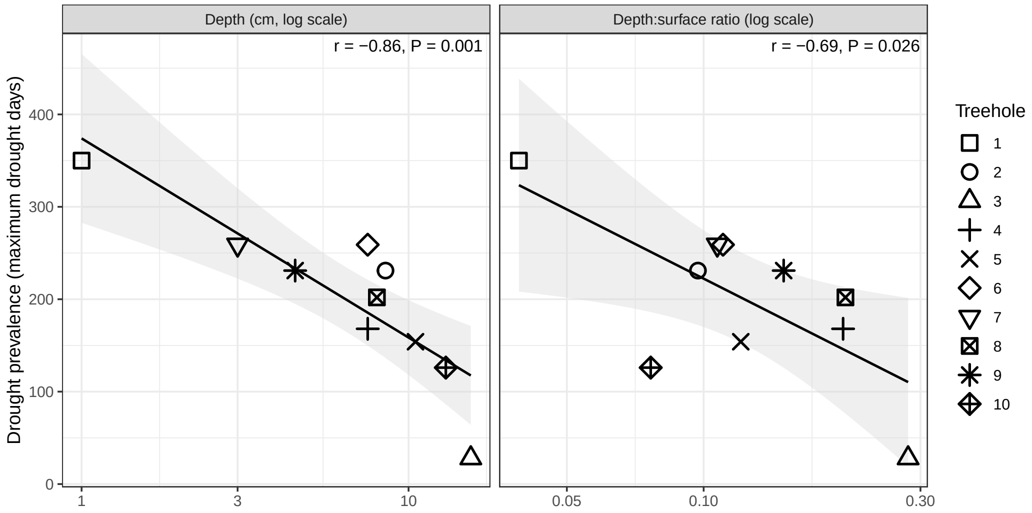


Figure 9: **Drought prevalence as a function of tree-hole depth and depth-to-surface ratio.** Each point is a tree-hole; drought prevalence is the maximum number of days that the tree-hole remained dry during the survey. Lines are least-squares fits with 95% confidence bands, and $r$ is the Pearson correlation of drought prevalence with the log-transformed variable ($n = 10$).

**Network of metacommunity M7.** The distributions of SparCC scores of some metacommunities showed tails departing significantly from the mean (scores $\gtrsim 0.5$ in main-text Fig. 5B), which represent potential candidates for ecological interactions. We examined the network of M7, which comprised two tree-holes (TH1 and TH7) and showed the strongest evidence of homogeneous selection (yellow cells in main-text Fig. 4A). Tree-hole-associated taxa had markedly different abundances in each tree-hole (main-text Fig. 5D), although all of them were present in both, and mapping this association onto the network showed that the two groups of the network corresponded to the taxa associated with each tree-hole (node colours in main-text Fig. 5C). Networks inferred independently for TH1 and TH7 were largely dissimilar, and only one small subnetwork had links conserved in both tree-holes, dominated by positive associations between ASVs from different taxonomic classes.

### Supplementary Figures

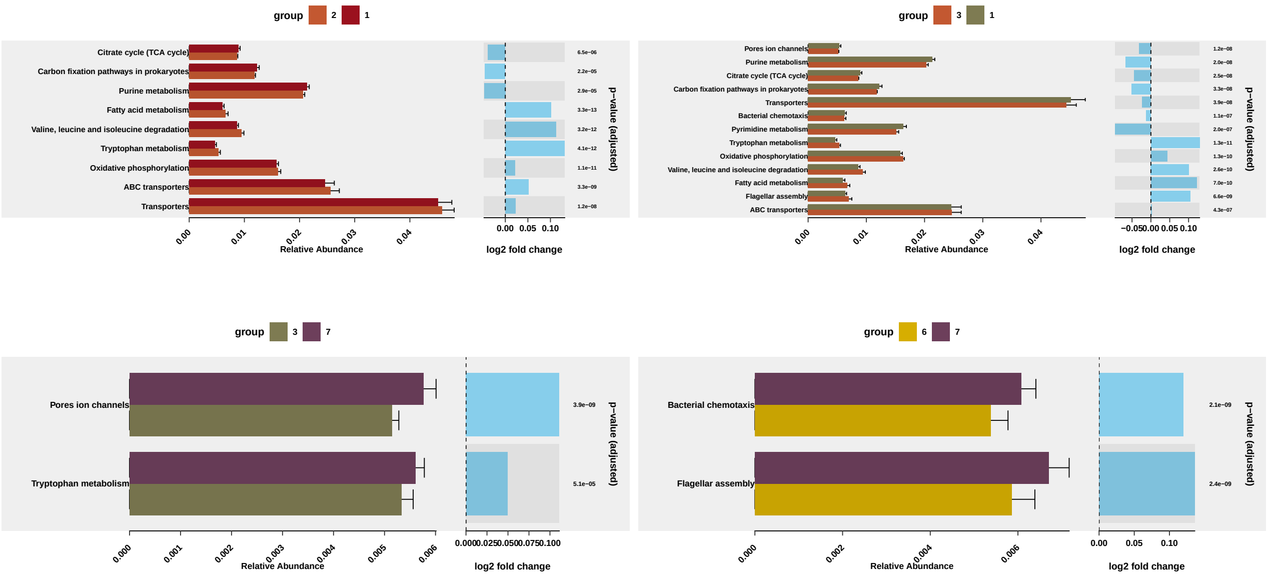


Figure 10: **Differences in the mean proportions of selected PICRUSt pathways**. Relative abundances of pathways significantly different between metacommunities differentiated by variable selection (top row) or dispersal limitation (bottom row). Variable selection leads to more significant pathways with higher effect sizes.

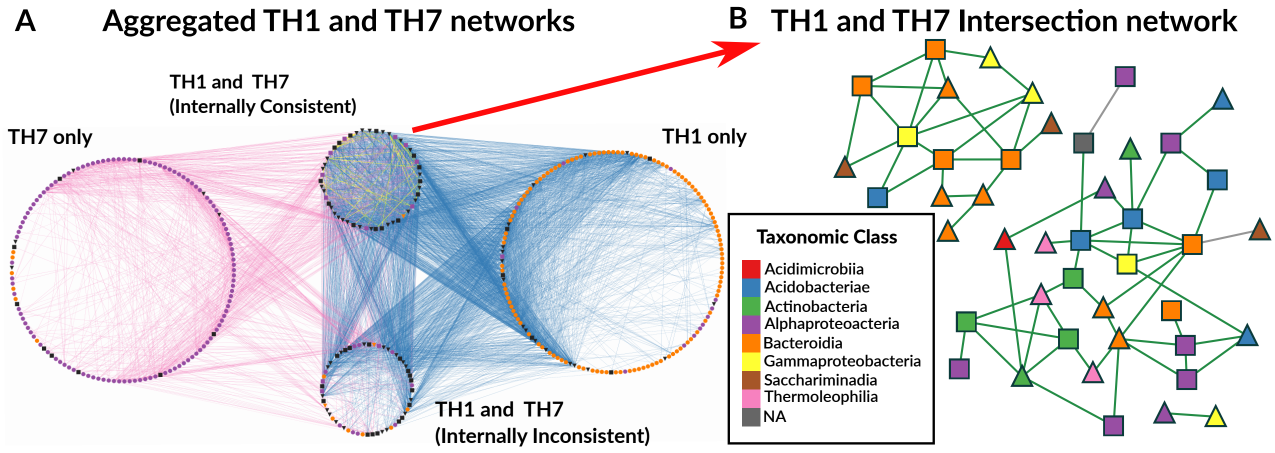


Figure 11: **Association networks.** (A) Network of selected ASVs grouped by functionInk guilds on each TH (exclusive of one TH) and those with links on both TH but that may have the same links (internally consistent) or not (inconsistent). Links denote TH where the interaction is found (blue: TH1, pink: TH7, yellow: both). Node colors denote rank-abundance selection (orange: TH1, purple: TH7, black: no preference). Node shapes denote IndVal selection (triangles: TH1, squares: TH7, circles: unselected). (B) Network of shared co-occurrence links from panel A. Node colors indicate taxonomic order; shapes denote combined IndVal classication (triangles: TH1, squares:TH7). Links are green (positive agreement) or grey (sign discrepancy).
